# Drosophila Amus and Bin3 methylases functionally replace mammalian MePCE for capping and the stabilization of U6 and 7SK snRNAs

**DOI:** 10.1101/2023.07.08.548165

**Authors:** Qiu Peng, Yiqing Wang, Ying Xiao, Hua Chang, Shishi Luo, Danling Wang, Yikang S. Rong

## Abstract

U6 and 7SK snRNAs process a 5’ cap, believed to be essential for their stability and maintained by mammalian MePCE or Drosophila Bin3 enzymes. Although loss of either protein results in 7SK instability, loss of neither is associated with U6 reduction. Their yeast homolog is also not required for U6 stability, casting further doubts on the function of capping U6. Here we show that the Drosophila Amus protein, homologous to both Bin3 and MePCE, is essential for U6 but not 7SK stability. A full function of Amus is required for Drosophila development, and that rests primarily on Amus’s methylase activity. Remarkably, the loss of U6 is rescued by the expression of an Amus-MePCE hybrid protein harboring the methyltransferase domain from MePCE, highlighting the conserved function of the two proteins as the U6 capping enzyme. Our new investigations in human cells establish a dependence of both U6 and 7SK stability on MePCE, resolving a long-standing uncertainty. While uncovering an interesting division of labor of Bin3/MePCE/Amus proteins, we discovered a “Bin3-Box” domain present only in enzymes associated with 7SK regulation. Targeted mutagenesis in Drosophila confirmed its importance for Bin3 function, revealing a possible conserved element in 7SK but not U6 biology.

## Introduction

RNA molecules are often modified at their 5’ extremity (the 5’ cap) with distinct functions *in vivo*. For examples, among RNAs produced by RNA polymerase II (Pol II), mRNAs are 5’ capped with an m^7^G, which is essential for their stability and translation (reviewed in Ramanathan et al. 2016). In addition, a class of small nuclear RNAs (snRNA) essential for pre-mRNA splicing (spliceosomal snRNAs), which includes U1, U2, U4 and U5, carries a tri-methylated cap (TMG) catalyzed by the Tgs1 enzyme (reviewed in Matera et al. 2007; Fisher et al. 2011). Results from *ex vivo* studies in vertebrate cells suggest that TMG is essential for the nuclear transport of snRNAs after their cytoplasmic assembly. However, genetic studies in yeast (e.g. Schwer et al. 2011) and flies (Cheng et al. 2021) contradict the notion that this is a fundamental function of TMG, and suggested other possibilities.

Pol III transcribed U6 spliceosomal snRNA (Singh and Reddy 1989) and the 7SK snRNA (Shumyatsky et al. 1990) have a unique 5’ cap in which the γ-phosphate of the guanosine triphosphate is methylated (γ-phosphate methylation). *Ex vivo* studies using synthesized RNA with or without this cap, and introduced into Xenopus eggs, led to the conclusions that this cap is essential for RNA stability (Shumyatsky et al. 1993), and that it prevents target RNAs from being translated (Chen et al. 2000), similar to what was additionally proposed for the TMG cap (Darzynkiewicz et al. 1988). Consistently, partial loss of the mammalian enzyme MePCE (Jeronimo et al. 2007), which catalyzes the capping reaction for 7SK *in vitro* (Yang et al. 2019), and the complete loss of the Drosophila MePCE-homologous Bin3 protein resulted in 7SK instability (Singh et al. 2011).

Molecularly, 7SK and U6 snRNAs share a similar stem loop structure at the 5’ end, which is essential for substrate recognition by the MePCE enzyme (Singh et al. 1990; Shumyatsky et al. 1994; Yang et al. 2019). Using a recombinant methylase domain of MePCE, it has been demonstrated that the enzyme transfers a methyl group from the SAM donor to both 7SK and U6 *in vitro* (Yang et al. 2019). Despite these results suggesting MePCE is the capping enzyme for U6, neither the partial loss of MePCE in mammalian cells nor the complete loss of Bin3 in Drosophila affects cellular U6 levels (Jeronimo et al. 2007; Singh et al. 2011). More recently, two groups carried out independent studies of the Bin3/MePCE homologous Bmc1 protein in *S. pombe*, an organism that lacks 7SK but depends on U6 for pre-mRNA splicing (Páez-Moscoso et al. 2022; Porat et al. 2022). Although U6 association with Bmc1 has been established in one study, both groups concluded that the loss of Bmc1 has no effect on U6 level. Therefore, after the discovery that U6 snRNA in many organisms has a γ-phosphate methylated cap over 30 years ago (Singh and Reddy 1989), ambiguity remains concerning the biological function of the U6 cap and the methyltransferase *in vivo*.

Here we characterized mutations in the gene encoding the Amus protein in Drosophila, which shares a methylase domain highly homologous to those in MePCE, Bin3 and Bmc1. Our results suggest that Amus is essential for U6 but not 7SK stability *in vivo*, and that this function depends on it being a methyltransferase. We re-investigated the effect of MePCE reduction in human cells, and obtained support that MePCE maintains U6 stability there as well. Remarkably, an Amus-MePCE hybrid protein with the methylase domain of MePCE is sufficient to restore U6 level in Drosophila *amus* mutants, showcasing the functional conservation of the two proteins. In addition, our phylogenetic study reveals two evolutionary trajectories of the more ancient U6 and the younger 7SK capping functions. We characterized a novel Bin3-Box motif, associated with the 7SK enzymes, highlighting a potential evolutionary driver for Bin3/MePCE/Amus divergence.

## Results

### Amus is a Bin3/MePCE homologous protein essential for animal development

There are three proteins annotated in flybase as homologous to mammalian MePCE enzyme: the original bicoid-interacting protein 3 (Bin3) identified by the Hanes group (Zhu and Hanes 2000), and the poorly characterized CG1239 and CG11342 proteins (Figure S1). CG11342 is more homologous to the recently identified tRNA capping methylase BCDIN3D (Figure S2) that might also affect the processing of miRNAs (Xhemalce et al. 2012; Martinez et al. 2017). The CG1239 protein has a well conserved methylase domain belonging to the Bin3/MePCE-type methylases (Figure S1). Although Drosophila Bin3 contains over a thousand residues and human MePCE close to seven hundred, CG1239 is significantly smaller and likely contains a single methylase domain, a situation similar to its *pombe* homolog Bmc1 (Figure S1). We rename CG1239 as <u>A</u>nother <u>M</u>ethylase of <u>U</u> <u>S</u>ix (AMUS), based on our results presented below.

Using CRIPSR/Cas9-mediated mutagenesis, we generated two mutations of *amus* (Figure 1A). The *amus^A-1-3^* allele carries a 10 bp deletion within the coding region leading to a frame shift and a premature translational STOP after the 97th residue. The *amus^1-1^* allele carries a mutation that changes the L^98^DV residues to I^98^FK. Due to the early STOP in *amus^A-1-3^*, we predict that it represents a severe or complete loss-of-function allele, while *amus^1-1^* a partial loss-of-function allele owing to the potential production of a full length but altered Amus protein. Consistently, animals homozygous for *amus^A-1-3^* (*amus^A-1-3/A-1-3^*), or heterozygous for *amus^A-1-3^* and a chromosomal deficiency (*df*) of the *amus* region (*amus^A-1-3/df^* trans-heterozygotes), die as second instar larvae, while animals that are *amus^1-1/A-1-3^*, or *amus^1-1/df^* survive to become sterile adults (Figure 1A). We constructed a rescuing transgene that carries a genomic fragment from the *amus* locus with its endogenous regulatory elements for transcription. Based on this transgene, we inserted a GFP-encoding fragment to the C-terminus of Amus coding region (*amus^gfp^*, Figure 1A). Both transgenes (with or without the GFP tag) rescue the early lethal or sterile phenotypes of *amus* mutants, further confirming that the phenotypes are due to the loss of Amus function.

**Figure 1.**
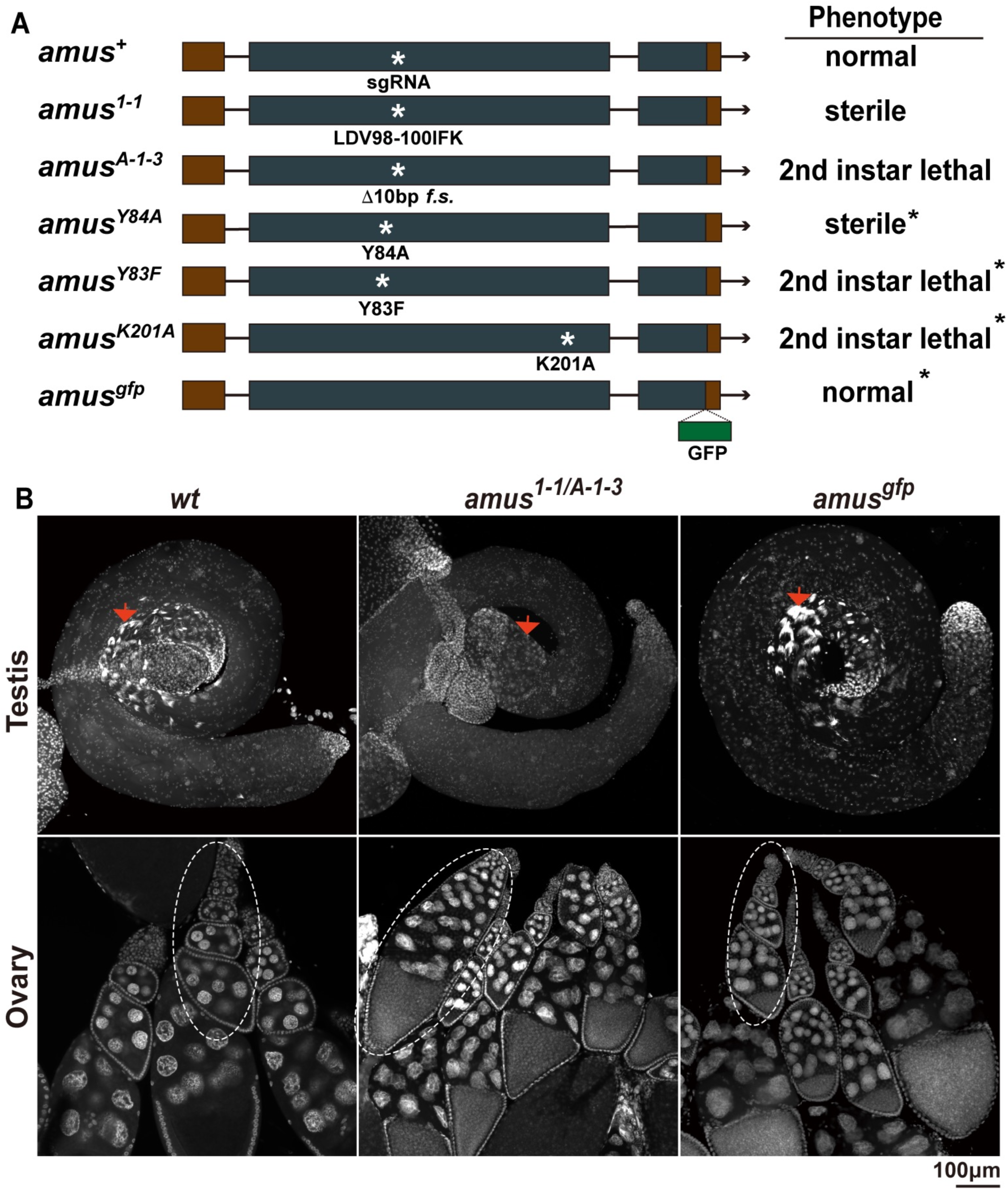
The mutations in *amus* and their phenotypes. **A.** Schematics showing the *amus* alleles. The allele names are list to the left of the gene diagrams. Rectangular boxes represent exons, with liver and blue colors representing non-coding and coding regions respectively. The arrow indicates the direction of transcription. The approximate positions of the sgRNA-targeting region and mutations generated in this study are indicated as white stars in the diagrams, with the amino acid residue changes indicated under. f.s.: frame shift. The phenotype of each allele is listed to the right of the diagrams. Asterisks represent cases in which the phenotypes were derived from animals that were also *amus* mutant (*amus^A-1-3/df^*). **B.** DAPI staining of testes and ovaries. The genotypes are listed at the top of the images. The arrows indicate areas in the testes where sperm bundles are visible in *wt* and *amus^gfp^*-rescued flies but not in *amus^1-1^*-mutant testes. In ovaries, a string of ovarioles is outlined with an oval, highlighting egg chamber fusions in the mutant but not wildtype ovaries.

Again based on rescuing transgenes, we constructed three additional mutations each of which changes a single conserved amino acid predicted to be critical for Amus’s methyltransferase function (Figures 1A and S1). The Y83F mutation changes a conserved Tyr residue, whose MePCE equivalent (Y421) is critical for interacting with the RNA substrate (Yang et al. 2019). The K201A mutation changes a MePCE K585 equivalent residue that is critical for both SAM and substrate binding (Yang et al. 2019). The introduction of these two mutant transgenes (*amus^Y83F^* and *amus^K201A^*) into either *amus^A-1-3/A-1-3^* or *amus^A-1-3/df^* background did not rescue of the early lethal phenotype. Although these results strongly suggest that the methyltransferase activity is essential for Amus function, we cannot rule out that both mutations destabilize the Amus protein as we have not been able to reliably detect the endogenous Amus protein on Western blots. We generated a third point mutation of Y84A, which alters another conserved Tyr residue in the area where MePCE interacts with its RNA substrate. Interestingly, introduction of this mutation to the *amus^A-1-3/A-1-3^*background resulted in development of the mutant animals into sterile adults, a phenotype similar to that caused by the *amus^1-1^* mutation.

We investigated the possible causes for the sterile phenotype of *amus^1-1^*and *amus^Y84A^* (full genotype *[amus^Y84A^]; amus^A-1-3^/amus^A-1-3^*) individuals by performing DAPI staining for DNA in ovaries and testes. As shown in Figure 1B, *amus*-mutant ovaries have significant development so that mature egg chambers are present.

Interestingly more than 25% of the ovarioles studied (n=47) display fused egg chambers in which multiple ones in a single ovariole are connected without the presence of follicle cells normally separating them (Figure 1B). Fused egg chambers have been previously observed in other mutations (e.g., Baksa et al. 2002; McGregor et al. 2002), suggesting distinct signaling pathway(s) are disrupted in *amus* mutant ovaries. However, this disruption could not have been the sole cause for female sterility as normal looking egg chambers are still present in the mutant. The cause of male sterility seems more straightforward. As shown in Figure 1B, *amus*-mutant testes lack post-meiotic development displaying the classic “meiotic arrest” phenotype (for reviews on the phenotype see Fuller 1998; White-Cooper 2010).

Therefore, the normal function of Amus is essential for animal development. Its complete loss or the specific loss of its methyltransferase function leads to early lethality while its partial loss disrupts germline development in both sexes.

### Amus is a nuclear protein that interacts with and controls the level of U6

We constructed an *amus^gfp^* strain in which the only functional Amus protein is produced from the transgene with a tagged *amus* coding region (complete genotype: *[amus^gfp^] amus^A-1-3^/[amus^gfp^] amus^A-1-3^*), and used GFP fluorescence as an indicator for Amus localization in cells. As shown in Figures 2A and S3A, Amus^GFP^ is highly enriched in the nucleus in cells from a variety of different tissues. This nuclear localization of Amus is consistent with the localization of U6 snRNA, one of its potential substrates (Figure S3B). Again, utilizing the GFP tag, we investigated whether Amus interacts with U6 by immunoprecipitating the Amus^GFP^ protein followed by Northern blot detection of U6. Results shown in Figure 2B reveal that U6 is present in GFP-IP using embryonic extracts made from *amus^gfp^*but not untagged *wt* animals, suggesting that Amus interacts with U6 *in vivo*.

**Figure 2.**
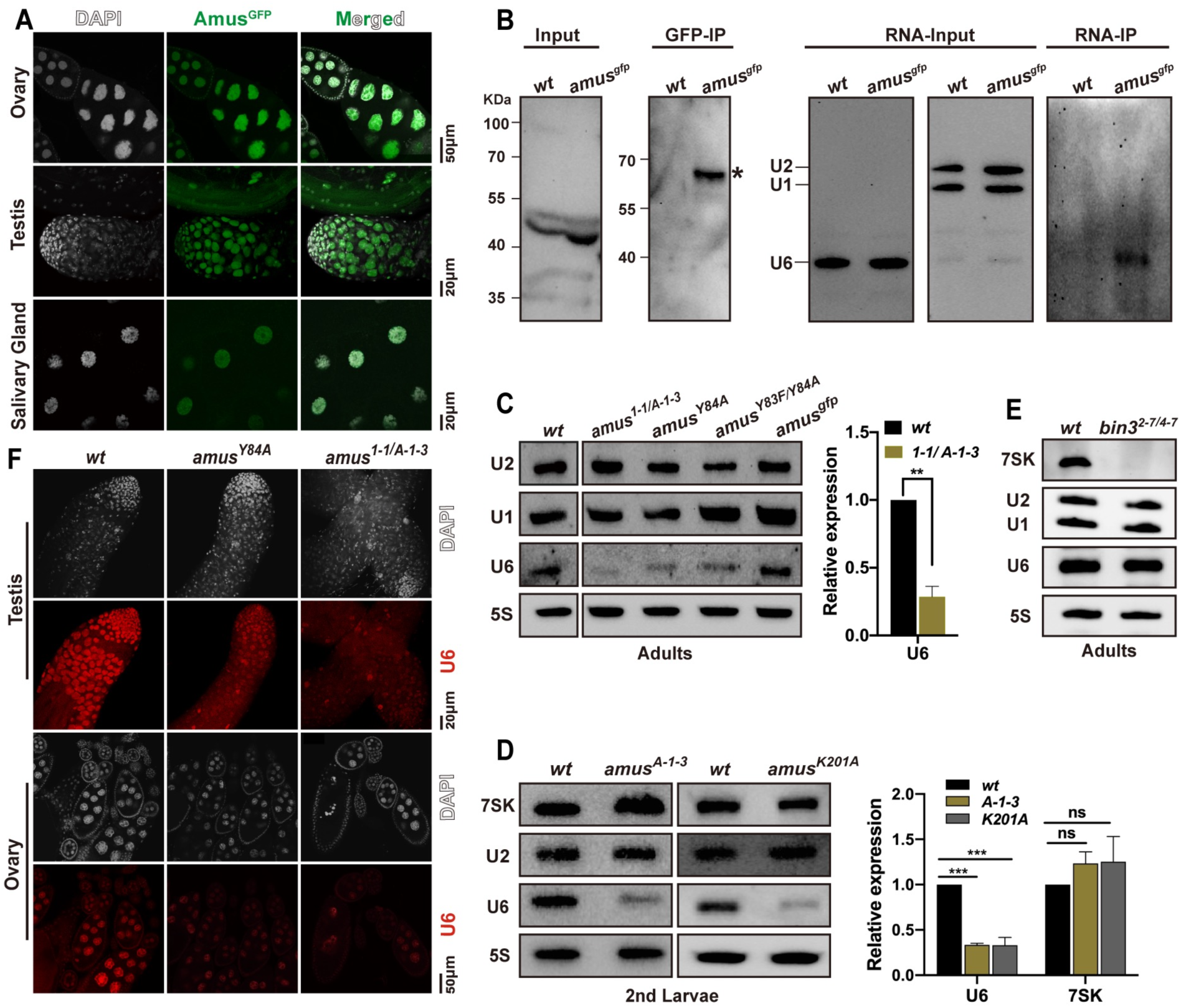
Amus is a nuclear protein essential for maintaining U6 level. **A.** Amus^GFP^ localization. GFP signals are in green, DNA in white. The examined tissues are ovaries and testes from adults and salivary glands from third instar larvae. More images can be found in Figure S3A. **B**. Testing U6-Amus interaction by RNA-IP. Two Western blots were shown to the left for the detection of Amus in embryonic extracts. The molecular marker positions are indicated to the left. The predicted molecular weights of Amus and Amus^GFP^ are approximately 38KDa and 68KDa respectively. In “Input”, only non-specific bands are visible. In “GFP-IP”, the asterisk labels the predicted Amus^GFP^ position. Three Northern blots are shown to the right for the detection of U6 with U1 and U2 serving as loading controls. In “RNA-IP”, U6 is detected only in the Amus^GFP^ IP sample. **C**. Northern blot analyses of U6 levels in *amus* mutant adults. Genotypes are listed on top and RNA species detected to the left. Quantification of U6 signals from three biological replicates are given in the chart with expression levels normalized against that of 5S rRNA. **D**. Northern blot analyses of U6 levels in *amus* mutant larvae. Genotypes are listed on top. Quantification of U6 signals from three biological replica are given in the chart with expression levels normalized against that of 5S rRNA. **E**. Northern blot analyses *bin3* mutant adults. Genotypes are listed on top. **F**. RNA FISH analyses of U6 in testis and ovary. DNA signals are in white and U6 signals in red. More images are shown Figure S3B. Genotypes are listed to the top. The full genotypes for this figure are: *amus^gfp^* (*[amus^gfp^] amus^A-1-3^*/*amus^df^*); *amus^Y84A^* (*[amus^Y84A^] amus^A-1-3^*/*amus^df^*); *amus^Y83F/Y84A^*(*[amus^Y83F^] amus^A-1-3^*/*[amus^Y84A^] amus^A-1-3^*); *amus^K201A^* (*[amus^K201A^] amus^A-1-3^*/*amus^df^*). **: *p*<0.01; ***: *p*<0.001.

Results from *ex vivo* studies from Xenopus suggest that the 5’ cap of U6 helps maintain its stability. If Amus were the U6 capping enzyme, we would expect a reduction of U6 level in *amus*-mutant cells. We used two methods to determine the level of U6: Northern blot analysis on total RNA and whole amount RNA FISH in germline tissues. In Northern blot analyses, the steady state level of U6 is significantly reduced in both *amus^1-1/df^* adults, and *amus^A-1-3/df^* second instar larvae, and this reduction can be rescued by the *amus^gfp^* transgene but not ones with disrupted methyltransferase activities (Figure 2C, D). Interestingly, U6 level remains unchanged in *bin3* mutants while that of 7SK is significantly reduced confirming prior results (Figure 2E). However, the level of 7SK remain unchanged in *amus* mutants (Figure 2C, D).

We also investigated whether loss of Amus affects the levels of other small RNAs. This was prompted by our earlier hypothesis that Amus is not the tRNA capping enzyme, and by prior results suggesting Amus/CG1239 is important for miRNA biogenesis (Zhu et al. 2019). Using Northern blot and RT-qPCR assays, we showed that the levels of various tRNA and miRNAs are not affected by *amus* mutations (Figure S4A, B, C). The *pombe* Bmc1 protein has been shown to be important for telomere length homeostasis by interacting with the telomerase RNA substrate (Páez-Moscoso et al. 2022; Porat et al. 2022), we therefore measured telomeric transcription in *amus* mutants but did not observe any significant change (Figure S4D). Therefore, loss of Amus seems to specifically affect the steady state level of U6 snRNA.

We conducted whole mount RNA FISH analyses in testes and ovaries from wildtype, *amus^1-1/df^* and *amus^Y84A^* adults. Using a probe complement to U6 transcripts, we detected a visible reduction of U6 in mutant tissues while the remaining U6 signals are nuclear, suggesting that the loss of Amus likely affects the stability of U6 but not its localization (Figures 2F and S3C). An antisense probe was included as a negative control in the analyses to confirm probe specificity (Figure S3D).

An anticipated outcome from U6 reduction is a reduction of pre-mRNA splicing efficiency that could result in intron retention. We previously showed that a *tgs1* hypomorphic mutation defective in synthesizing the TMG cap for the other spliceosomal snRNAs results in wide-spread intron retention in testis (Cheng et al. 2020). Using the same PCR-based approach, we detected splicing defects in testis of the hypomorphic *amus^1-1/df^* mutants (Figure S5). The hypomorphic *tgs1* mutations results in male sterility with a “meiotic arrest” phenotype highly similar to that observed in *amus^1-1/df^* testis (Figure 1B). Similarly, the pleiotropic effect from splicing dysfunction could explain sterility for both *tgs1* and *amus* mutant males.

In summary, the Amus protein is localized to the nucleus where it interacts with U6 snRNA. Loss of Amus leads to a reduction of U6 and causes defects in pre-mRNA splicing, and the methyltransferase activity of Amus is essential for its functions in development.

### Amus’s methyltransferase function is replaceable by that of MePCE

Our results thus far are consistent with that Amus caps U6 to maintain RNA stability, a function assigned to the mammalian MePCE based on *in vitro* methylation results. The two proteins share significant homology between their methyltransferase domains (Figure S1), which suggests that the mammalian methyltransferase might be functional in flies as their suspected substrate of U6 is highly conserved. As a way to gather *in vivo* evidence supporting MePCE is the U6 capping enzyme, we tested the above hypothesis taking advantage of the Drosophila *amus* mutants. Since the mammalian protein has to be expressed in Drosophila from an intron-less transgene, we first investigated the proper expression condition using an *amus* cDNA clone as a control. We placed it under the control of *UAS* elements so to drive its expression by the Gal4 activator. We discovered that driving *amus*-cDNA expression with the ubiquitous actin5C-Gal4 driver rescued the lethality of *amus^A-1-3^*homozygous and *amus^A-1-3/df^* trans-heterozygous animals. Moreover, the rescued viability is accompanied by a restoration of the level of U6 on Northern blots (Figure 3C).

**Figure 3.**
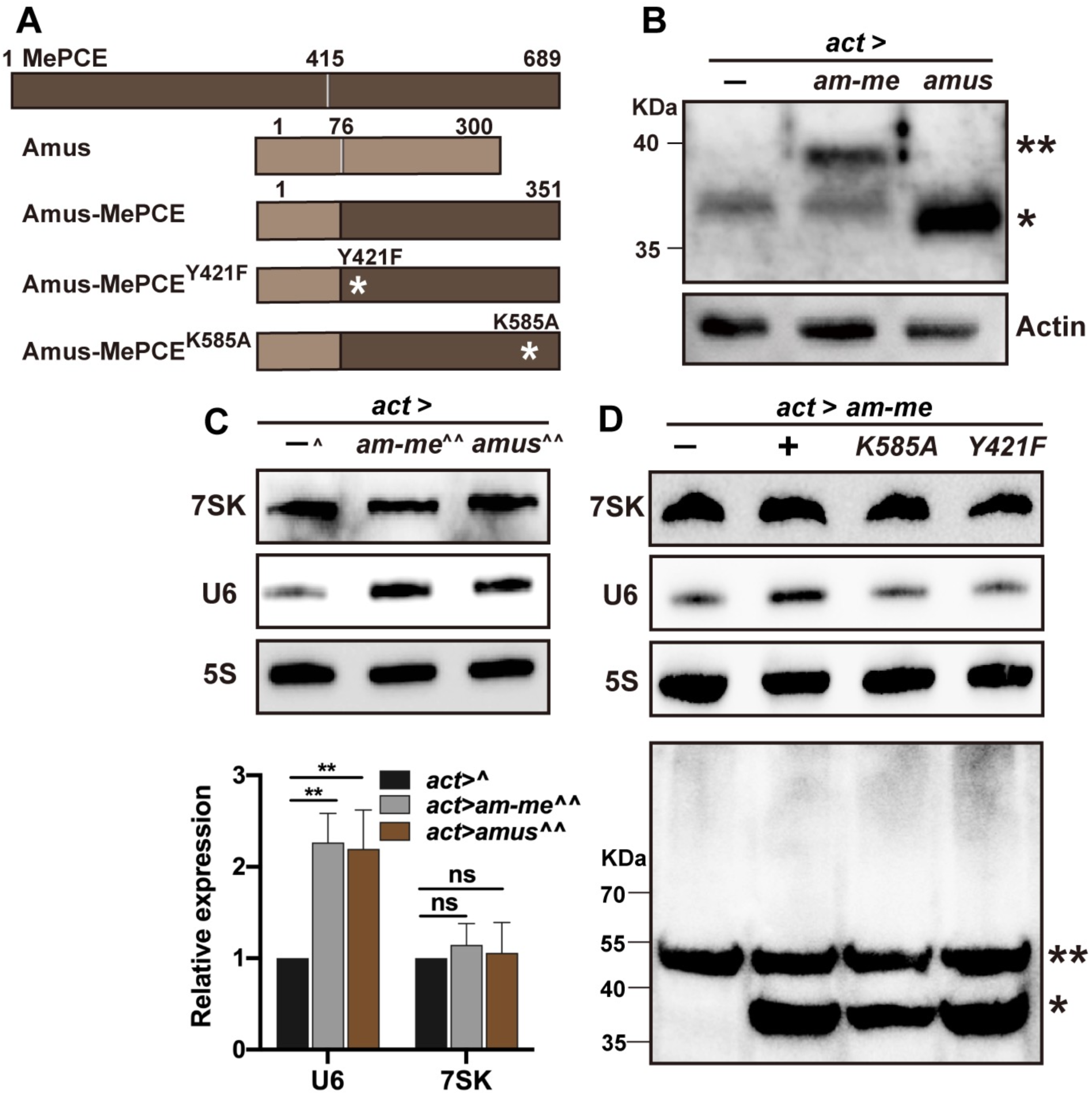
The methylase function of Amus can be replaced with that of MePCE. **A**. Diagrams of Amus, MePCE and Amus-MePCE hybrid proteins. The rectangular boxes represent proteins with Amus and MePCE drawn in different colors. Amino acid residue numbers are indicated on top of the boxes. In the hybrid protein, residues 415 to 689 from MePCE replaces 77 to 300 of Amus. The star represents the approximate position of single amino acid mutations with MePCE designations. **B**. Western blot showing the production of the overproduced proteins. Total extracts were produced from wildtype adults that carried the actin5C driver (*act>*) either alone (-) or with Amus (*amus*) or Amus-MePCE (*am-me*) expression constructs. The double star marks the position of Amus-MePCE. Since Amus-MePCE retains one quarter of the Amus antigen, its signal on Western blots is expected to be significantly weaker than that of Amus. The single star marks a non-specific band. **C**. Northern blot detection of U6 level in flies. Total RNA was extracted from animals that carried the actin5C driver (*act>*) either alone (-) or with Amus (*amus*) or Amus-MePCE (*am-me*) expression constructs. The *act>-* animals (^) are also heterozygous at the endogenous *amus* locus (*amus^df/+^*). The *act>amus* and *act>am-me* animals (^^) are mutant at the endogenous locus (*amus^A-1-3/df^*). Quantification of U6 level from three biological replicates is shown under with expression levels normalized against that of 5S rRNA. Overexpression of either Amus or Amus-MePCE results in elevation of U6. *: *p*<0.05; **: *p*<0.01. **D**. Effects on U6 from Amus-MePCE overexpression in Drosophila S2 cells. At the top is Northern blot results where we used actin5C-Gal4 to drive expression of Amus-MePCE and its methylase dead derivatives of K585A and Y421F. Below is Western blotting with anti-MePCE showing the expression levels of Amus-MePCE and derivatives, marked with a single star. The double star marks Tubulin as a loading control.

The C terminal methyltransferase domain of MePCE is able to methylate U6 *in vitro*. We thus replaced the homologous domain in Amus with this half of MePCE leaving the first 76 Amus residues in place (Figure 3A, B). A nuclear localization signal is predicted within this remaining Amus fragment, potentially preserving the ability of the Amus-MePCE hybrid protein to be properly localized. Using the same actin5C-Gal4 driver, we delivered Amus-MePCE to *amus^A-1-3^* homozygous or *amus^A-1-3/df^* trans-heterozygous animals. We showed earlier that loss of Amus results in death at the second instar larval stage. Remarkably, the expression of Amus-MePCE prolongs the survival of these animals: all reaching the pupal stage (an extra four days of development). More importantly, the ubiquitous expression of the hybrid protein restores the level of U6 (Figure 3C). We were led to conclude that Amus-MePCE is proficient in maintaining the level of U6, possibly as its capping enzyme *in vivo*. This conclusion is supported by results from additional experiments in which we individually expressed two Amus-MePCE variants that harbor mutations of critical amino acid residues essential for its methyltransferase activity (Figure 3A). None of these variants were able to rescue the viability of *amus* loss-of-function mutants beyond the second instar stage. Therefore, the molecular function of Amus and MePCE as the methyltransferase for U6 snRNA is similar.

We noticed that the U6 level in flies overexpressing Amus or Amus-MePCE reaches above that of wildtype animals (Figure 3C). We noted that the endogenous U6 level also elevates above the normal level in S2 cells overexpressing the Amus-MePCE hybrid protein, and that this effect depends on MePCE’s methylase activity (Figure 3D). We do not consider that this overproduction of U6 is likely responsible for the failure of the Amus-MePCE expressing animals to reach adulthood since Amus overexpressing animals survive normally. Instead, we propose that the hybrid protein might not have been able to fully carry out Amus’s other and perhaps U6-indepednent functions in Drosophila.

### MEPCE is essential for U6 stability in mammalian cells

After having obtained evidence supporting that MePCE can be a U6 capping enzyme *in vivo*, we re-visited the issue of whether MePCE reduction in mammalian cells would result in U6 instability. We designed four different siRNAs targeting different regions of the *mepce* transcript (Figure 4A), and confirmed MePCE reduction by Western blotting. As shown in Figure 4B, all four siRNA treatments resulted in efficient reduction of MePCE in HEK293 cells, the same cell line used in the earlier study (Jeronimo et al. 2007). As a result, the level of the 7SK snRNA is greatly reduced as expected. Interestingly, the level of U6 is also reduced significantly (Figure 4C). We chose two efficient siRNA reagents and repeated the experiments in HeLa cells. Results shown in Figure 4D clearly suggest that U6 level drops upon MePCE reduction in this second cell line as well. Therefore, MePCE is required for maintaining the steady state levels of both 7SK and U6 in human cells, consistent with its role as the capping enzyme for both snRNAs.

**Figure 4.**
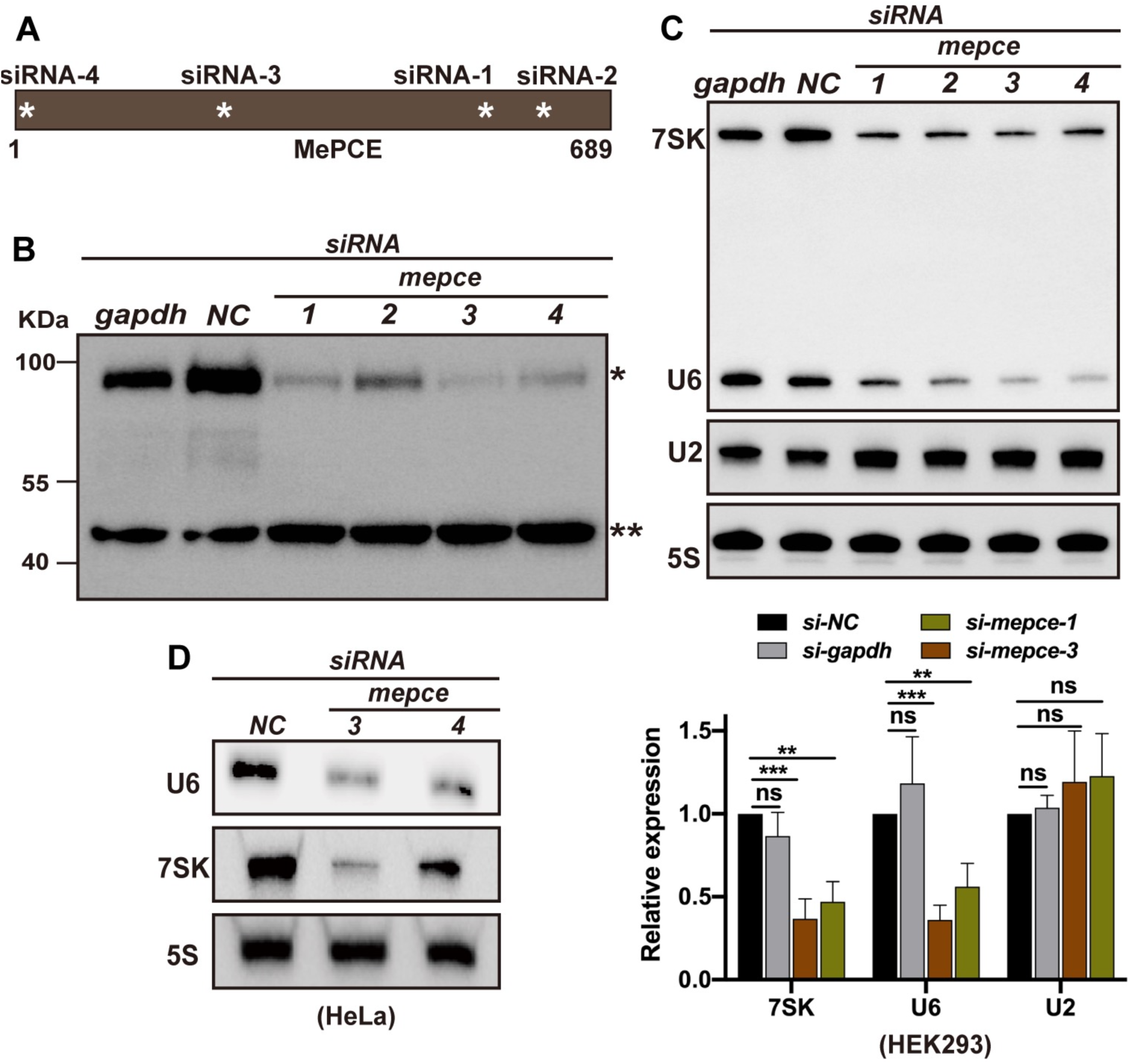
Human MePCE is essential for U6 stability. **A.** Diagram showing the approximate position of siRNA targeted regions of the MePCE. Each position is marked with a white star. The numbers below are amino acid residue numbers of MePCE. **B**. Western blot verification of the effectiveness of MePCE knock-down in HEK293 cells. The siRNA targets are listed on top of the gel, with protein marker positions indicated to the left and target proteins: MePCE (*) and Actin (**) indicated to the right. Two control siRNAs are used: one against *gapdh* and one with randomized sequences (NC: negative control). **C**. Northern blot detection of U6 and 7SK in HEK293 cells. The siRNA targets are listed on top of the gel. U2 snRNA and 5S rRNA are used as controls. Quantification of U6 and 7SK levels are shown in the chart below with expression levels normalized against that of 5S rRNA. **: *p*<0.01; ***: *p*<0.001. ns: not significant. **D**. Northern blot detection of U6 and 7SK in HeLa cells. The siRNA targets are listed on top of the gel. 5S rRNA is used as a control.

### “Bin3-Box”: a conserved domain in 7SK regulating enzymes

Our results suggest an interesting evolutionary scenario in which the combined functions of Drosophila Bin3 and Amus are fulfilled by MePCE in mammals. To decipher the evolutionary events that might have led to this outcome, we conducted a sequence analysis of Bin3/MePCE/Amus-homologous proteins in eukaryotic organisms. Based on the phylogenetic analyses of 7SK and its binding partners in metazoan as a guide (Marz et al. 2009), we conducted BLAST searches using Drosophila Bin3, Human MePCE and Drosophila Amus as individual queries, and discovered orthologous proteins in many metazoan species. The Amus protein and its yeast homolog Bmc1 are the smallest of the three classes, and share significant homology to Bin3 and MePCE that is limited to the methyltransferase domain. The two larger proteins, Bin3 and MePCE, share homology beyond the methyltransferase domain. In particular, a stretch of about 70 residues is conserved amongst Bin3/MePCE proteins (Figure 5A, B). We name this domain the “Bin3-Box”, which is likely present only in Bin3/MePCE related proteins based on BLAST searches.

**Figure 5.**
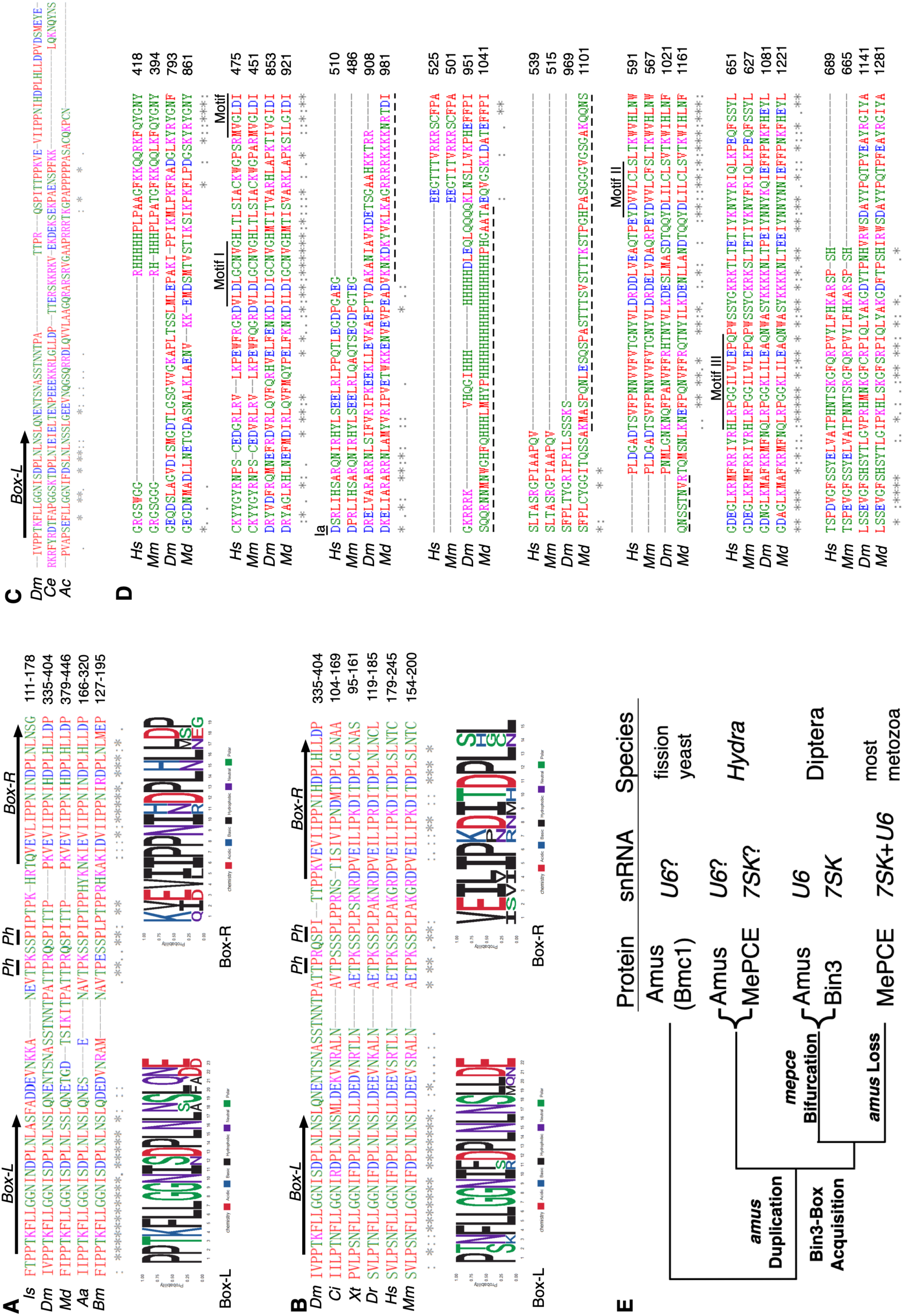
Sequence divergence reveals functional evolution of the Bin3/MePCE/Amus proteins. **A**. Bin3-Box from arthropod Bin3/MePCE proteins. At the top is the alignment of representative Bin3 proteins. Above the alignment, the approximate positions of the two tandemly repeated boxes (L: left; R: right) are denoted by arrows. The two conserved positions where potential phosphorylation of the Thr and Ser residues are denoted with “Ph”. At the bottom is the diagram showing amino acid distributions at different positions of the two boxes. Note that the short motif of “NI*DPL*L” is present and conserved in both boxes. Is: *Ixodes scapularis*, Dm: *Drosophila melanogaster*, Aa: *Aedes aegypti*, Bm: *Bombyx mori*. **B**. Bin3-Box from metazoan Bin3/MePCE proteins. Note that the right box is more degenerated when compared with the alignments shown in **A**. Ci: *Ciona intestinalis*, Xt: *Xenopus tropicalis*, Dr: *Danio rerio*, Hs: *Homo sapiens*, Mm: *Mus musculus*. **C**. A “half” version of the Bin3-Box. The two species where a Bin3-Box with only the left copy is clearly identifiable are used to align with Bin3 from *Drosophila melanogaster*. Ce: *Caenorhabditis elegans*, Ac: *Anolis carolinensis*. **D**. Bifurcation of the Dipteran Bin3 methyltransferase domain. Sequence alignments of the methyltransferase domain are shown with structural “Motifs” indicated with solid lines. The larger insertions between Motif Ia and Motif II in Dipteran proteins are indicated by dotted lines. **E**. A model highlighting the evolutionary events during Bin3/MePCE/Amus divergence. The proteins, their potential snRNA substrates and representative species are listed to the right of the tree. Four crucial events (*amus* duplication, Bin3-Box acquisition by one of the two *amus* copies, Bifurcation of the methylase domain in the Diptera lineage, and frequent loss of *amus* throughout metazoan evolution), and their proposed timing are indicated.

Upon closer inspection of Bin3-Boxes, we uncovered a few of its interesting features. First, a Bin3-Box appears to be made of two tandem peptide repeats. This feature is prominent when arthropod proteins are aligned (Figure 5A), showing a repetition of the NL*DPL*L motif. While the left copy (N-terminal) is highly conserved amongst Bin3/MePCE homologs shown in Figure 5B, the homology amongst the right ones is more degenerated even within the highly conserved “DPLNL” motif.

Secondly, the two repeated copies flank a short peptide with “TP” and “SP” at conserved positions (Figure 5A and B), both positions are predicted to have a high potential to be phosphorylated. Thirdly, we noticed instances in which the repeated copy at the right is highly degenerated including the “DPLNL” signature so that only the left copy can be identified. The best examples are found in Bin3/MePCE from the model worm *Caenorhabditis elegans* and the lizard *Anolis carolinensis* (Figure 5C). In these proteins with a half Bin3-Box, the TP/SP sites are also missing. Therefore, whether complete or truncated, Bin3-Box is present in Bin3/MePCE class of proteins, but not Amus/Bmc1-related ones. We suggest that this is related to Bin3/MePCE, but not Amus/Bmc1, being the 7SK capping enzyme.

### The acquisition of Bin3-Box and bifurcation of the methylase domain as potential drivers for the evolution

Within Dipteran insects, we observed widespread co-existence of both Bin3 and Amus homologs. Surprisingly, this division of labor does not appear to occur in insects outside of Dipteran species, or in most metazoans. As noted by Cosgrove et al. (2012) before, the methyltransferase domain of Drosophila Bin3, when compared with that of MePCE, is bifurcated by an insertion, between Motif Ia and Motif II, of about 50 residues highly enriched with charged residues (Figure 5D). A similar situation exists for Bin3 from the mosquito *Aedes aegypti*, and to a more extreme extent for Bin3 of the house fly *Musca domestica* in which two insertions amounting to more than 100 residues have occurred between Motifs Ia and II (Figure 5D). We speculate that these “extra” residues might have interfered with Bin3’s ability to target U6 snRNA.

We suggest that Amus is the “ancestral” gene product due to the presence of homologous proteins in the fission yeast (Bmc1), the model plant *Arabidopsis thaliana*, and ciliates (Table S1). It is therefore more logical to attribute the missing of Amus in most metazoans to gene loss rather than gain in Dipterans. Our proposition is supported by the discovery of Hydra as a second lineage with separate genes encoding a Bin3 and an Amus like proteins (Table S1).

We devise a speculative model in Figure 5E to help trace the evolution of Bin3/MePCE/Amus proteins and their functions. The ancestral function of Amus is to cap U6 snRNA. The birth of 7SK coincided with a duplication of the ancestral *amus* gene accompanied by the acquisition of the Bin3-Box by one of the copies, giving rise to the prototype of MePCE. The redundant Amus was later lost in most metazoans as MePCE is able to supply Amus’s function of protecting U6. In Dipteran, however, the bifurcation of the methyltransferase domain abolished Bin3’s ability to cap U6, necessitating the presence of Amus in these organisms.

### The Bin3-Box is essential for the *in vivo* function of Bin3

An important prediction of the above model is that the Bin3-Box is a functional element of Bin3-class proteins. To gather supporting evidence, we used CRISPR/Cas9 to specifically disrupt the Bin3-Box of Drosophila Bin3. This is possible because CRISPR/Cas9 induced small deletions do not necessarily result in frameshift mutations. We designed sgRNAs to the region of *bin3* encoding the Bin3-Box and was able to recover multiple *bin3* alleles carrying small in frame deletions (Figure 6A, S6). We chose three of the newly generated alleles to assay the functional importance of the Bin3-Box. *bins^20^* is a 54 amino acid residues deletion of almost the entire Bin3-Box; *bin3^63^* loses 6 amino acid residues of Bin3-Box, four of which are highly conserved residues (Figures 5A, B and 6A); and *bin3^68^* has a 4bp deletion accompanied by a frameshift mutation likely leading to a complete loss of Bin3 function.

**Figure 6.**
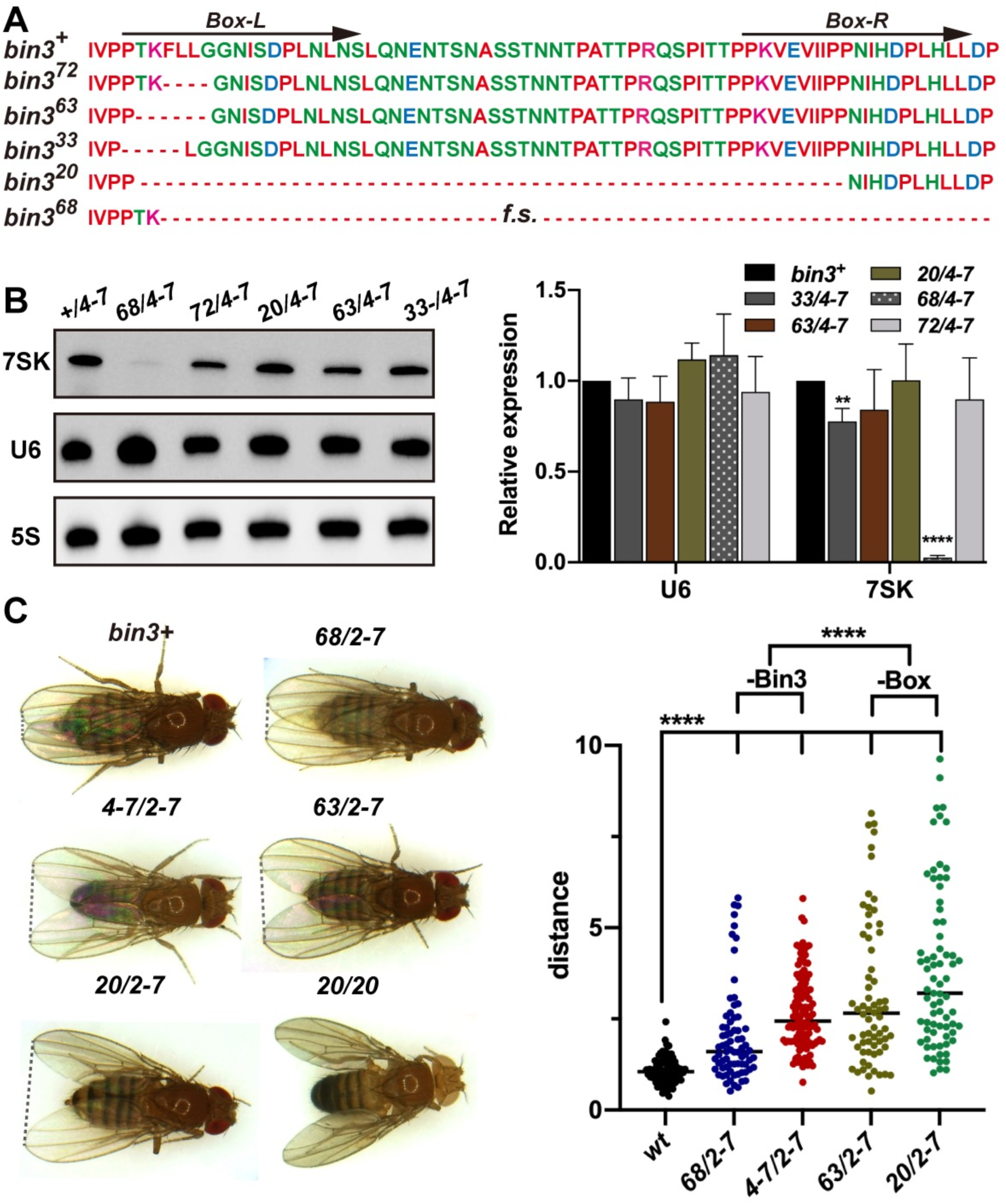
Bin3-Box is essential for Bin3 function in Drosophila. **A.** Amino acid sequences showing the *bin3* alleles. The wildtype sequence of the Bin3-Box is shown at the top. In the mutant alleles generated in this study, a deleted amino acid residue is indicated by a “-”. The *bin3^68^* allele is a frame shift (f.s.) mutation potentially losing the protein region C-terminal to the mutation. **B**. Northern blot analyses of *7SK* and U6 levels in *bin3* mutant adults. The *bin3* allelic combinations are shown at the top. Quantification of RNA levels from three biological replicates is shown in the chart with expression levels normalized over that of 5S rRNA. ****: *p*<0.0001. **: *p*<0.01. **C**. Representative pictures showing different wing postures from *bin3* adults. Allelic combinations are shown above the fly pictures. A dotted line connects the tips of the two wings and the distance was measured. Quantification of the measurement in different genotypes is given in the chart to the right. The differences of that distance were analyzed by t-tests using Prism and P values are calculated accordingly. ****: p<0.0001. A comparison was also made between flies carrying frame shift mutations of *bin3* likely missing the Bin3 protein (-Bin3) and those carrying in-frame deletion of the Bin3-Box only (-Box).

The newly generated *bin3* mutations, similar to the previous *bin3^2-7^* and *bin3^4-7^* alleles (Singh et al. 2011), support viability, and homozygous mutants can be maintained as stocks. Unlike the prior alleles of *bin3*, the newly generated *bin3* alleles do not result in a significant reduction of 7SK level except the frameshifted *bin3^68^* allele (Figure 6B). Therefore, the complete loss of the Bin3 methylase function is likely required for the loss of 7SK stability. These results would have suggested that Bin3-Box either does not carry any function or carries one that is not related to Bin3’s function in regulating 7SK function.

We noticed that homozygous adults of all newly generated *bin3* alleles share a consistent abnormality concerning the wing posture. As shown in Figure 6C and S7, these adults suffer a “held-out-and-down” wing phenotype in which the wings hold out laterally and down, instead of extending straight from the notum and partially crossing each other. We quantified the phenotype (Figure 6C) using flies that are trans-heterozygous for a *bin3* allele and the *bin3^2-7^* or *bin3^4-7^*deletion allele, minimizing the potential effects from second site mutations. It became clear that both *bin3* frameshift mutations and ones deleting only the Bin3-Box cause the wing posture phenotype.

Interestingly, the Bin3-Box affected alleles seem to have a stronger effect on wing posture. Remarkably, homozygous mutants of these alleles, in particular the Bin3-Box-deleted *bin3^20^* allele, have the strongest wing defects, suggesting that these alleles are semi-dominant in nature. This is possible as these mutations retain the ability to encode an altered Bin3 protein. Unfortunately, the lack an Bin3 antibody has prevented us from probing the presence of these defective proteins. Nevertheless, we conclude that Bin3-Box is an essential part of the Bin3 function even though it might not pertain to the maintenance of 7SK stability.

## Discussion

The U6 snRNA is one of the most conserved small RNA in nature and carries essential function in RNA splicing (for a review on U6 see Didychuk et al. 2018). Newly synthesized mouse U6 has a shorter half-life than those of other spliceosomal snRNAs (Sauterer et al. 1988). Evidence exists suggesting that the intra-cellular level of U6 is tightly controlled in human cells (Noonberg et al. 1996). It has been known for over thirty years that U6 snRNA from human and many other organisms is uniquely capped by methylation at its 5’ extremity and *ex vivo* studies led to the suggestion that the cap is required for U6 stability. However, RNAi knockdown of the presumed mammalian capping enzyme MePCE or deletion of its homologs in flies or yeast does not affect U6 level, casting doubts on the biological function of the cap on U6. Here we characterize MePCE and its Drosophila homolog Amus, and present genetic evidence for the first time supporting a protective role of the U6 cap in flies and men.

### Bin3/MePCE/Amus related proteins are important for U6 stability

The first attempt in identifying the U6 capping enzyme resulted in the purification of an activity able to bind and cap U6 *in vitro* (Shimba and Reddy 1994). Remarkably, the same activity efficiently caps 7SK, consistent with one of the major conclusions from our study that human MePCE caps both U6 and 7SK *in vivo*. We applied multiple siRNA agents targeting multiple regions of the *mepce* transcript, measured protein level to ensure effective MePCE reduction, and tested two different human cell lines for the effects of *mepce* KD. Our results are inconsistent with the long-held belief that MePCE is dispensable for U6 stability. We note that the original result suggesting otherwise was reported in 2007. Technology advances over the years might have allowed us and others to achieve more efficient MePCE elimination by RNAi. Unfortunately, the level of U6 snRNA was not monitored, to our knowledge, in any subsequent MePCE study. Nevertheless, the different effects on U6 stability from our and the 2007 studies suggest that 7SK stability is more susceptible to MePCE reduction than that of U6. U6 is in complex with a myriad of proteins and snRNAs forming multiple snRNPs (reviewed in Didychuk et al. 2018). This likely provides other protecting mechanisms in addition to the cap structure installed by MePCE. Such functional redundancy could also explain the normal steady state level of U6 in yeast *bmc1* mutants. MePCE and its homologs are not known components of any U6 snRNP. On the other hand, MePCE is an important component of the 7SK snRNP (Wassarman and Steitz 1991; Jeronimo et al. 2007; Nguyen et al. 2012), and known to play a protective role independent of its capping function (Xue et al. 2010), which potentially renders 7SK less resistant to MePCE loss.

The sensitivity of U6 to the loss of Amus in Drosophila is remarkable as even hypomorphic mutations that support full somatic development result in U6 reduction. We consider cell type or tissue specific effects as a potential explanation, as we imagine that somatic cells highly active in metabolism might require higher U6 turnover to support higher levels of transcription. Indeed, cell proliferation is limited as flies enter a growth phase during larval development, and *amus* loss-of-function mutation ceases growth during the second of the three instar stages, a stage when maternally deposited Amus protein and/or capped U6 RNA might run out. A thorough understanding of the physiological consequence from the loss of γ-monomethylated cap requires further efforts. We suggest that Drosophila offers an excellent genetic model for this dissection as hypomorphic *amus* mutations produce intermediate phenotypes that are well suited for genetic screens for enhancer/suppressor mutations.

### Protein-RNA interactions as a potential driver for Bin3/MePCE/Amus evolution

Our results uncovered an interesting evolutionary case of Sub-functionalization in which the activities of the mammalian MePCE enzyme are being fulfilled by two separate proteins in Drosophila. A single enzyme responsible for capping U6 and 7SK snRNAs might appear counterintuitive considering the very different evolutionary trajectories that the two RNA species are undertaking. While U6 is the most conserved snRNA in nature, 7SK is considered fast evolving so that its primary sequence, size or number are poorly conserved. Great efforts utilizing promoter sequence analyses and secondary structure predictions are needed for their identification in invertebrates and particularly so in arthropods (Gruber et al. 2008a, 2008b; Marz et al. 2009; Yazbeck et al. 2018). Therefore, the selective pressures placed on the single MePCE protein might dictate different rates of evolution at different parts of the protein. One could imagine that the methyltransferase domain is under more constrains while the rest of protein is likely more relaxed or even undergoing positive selection. Our demonstration of the ability of MePCE’s methylase domain to function in Drosophila is consistent with this hypothesis. Generally, our analyses are consistent with that the methyltransferase domain is conserved in animal, plant and fungal species while the rest of the protein is highly diverged with the dipteran Bin3 proteins being the extreme both in lengths and sequences.

Nevertheless, we identified an island of homology as the Bin3-Box. Bin3-Box, including its degenerative half version, is associated with Bin3/MePCE-homologous proteins throughout metazoan evolution. Its absence in homologs of Amus from Ciliates, Fungi and Plants but presence in Trichoplax (Table S1) leads us to suggest that it arose concomitantly with 7SK snRNA in animals, since evidence exists supporting the presence of 7SK in Trichoplax (Marz et al. 2009).

In Dipteran insects and likely Hydra, the capping functions of MePCE is fulfilled by Bin3 and Amus combined. Since the *amus* gene likely represents the ancestral copy, its absence as a stand-alone copy in most metazoan species would reflect its frequent losses. This is perhaps expected since MePCE (with a Bin3-Box) is capable of capping both snRNA species while Amus lacks the 7SK-protecting function. If so, what has resulted in the retention of Amus in Diptera? We and others has noticed the bifurcation of the methyltransferase domain happened to Dipteran Bin3 proteins. We speculate that the large insertion inside the methyltransferase domain abolishes Bin3’s ability to recognize U6, necessitating the retention of Amus in these organisms. However, the presence of both proteins in Hydra might need alternative explanations since Hydra’s Bin3/MePCE does not seem to be bifurcated.

The model presented in Figure 5E is likely over simplified. As the molecular functions of Bin3/MePCE/Amus are still being discovered, we have only started to understand the functional requirements driving protein evolution. For examples, evidence supports a role of Drosophila Bin3 in translational repression (Cosgrove et al. 2012). Moreover, Bmc1 is shown recently to have capping independent roles in telomere maintenance. Furthermore, one must not ignore the fact that budding yeast, planaria (Schmidtea) and chickens (Gallus) lack Bin3/MePCE/Amus all together even though a 7SK gene has been clearly identified in chicken (AJ890101, Egloff et al. 2006). How the snRNA-protecting functions are fulfilled in these organisms promises to be fascinating.

### Bin3-Box as a new functional module independent of RNA protection

The physiological effects of Bin3/MePCE loss have not been explored extensively with one publication focusing on defects in oogenesis and embryogenesis of Drosophila *bin3* mutants (Singh et al. 2011), one on its potential role in regulating tumor growth in Drosophila (Nagarkar et al. 2020), and one on its role during zebra fish development (Barboric et a. 2009). Interestingly, neuromuscular defects have been described for a young patient with a heterozygous *mepce* mutation (Schneeberger et al. 2019).

Using targeted mutagenesis, we investigated the function of the newly identified Bin3-box in Drosophila Bin3 protein. Unlike the complete loss of Bin3, loss of Bin3-box alone does not significantly impair 7SK level, suggesting that it has little or no role in maintaining the methylase function of Bin3. However, we identified defective wing posture as a novel physiological consequence shared by mutants that lose the Bin3 methylase or just the Bin3-box alone, suggesting that Bin3-Box is an essential part of Bin3 functions.

Defects in wing posture have been described for a large variety of mutant conditions in Drosophila, with underlining causes associated with muscle dysfunction, neuromuscular defects, redox abnormality and others (e.g., Zhang et al. 2001; Huang and Stern 2002; Anh et al. 2011; Johnstone et al. 2013; Lerch et al. 2020; Hinz et al. 2021). In a Ph.D. thesis deposited online (http://doi.org/10.25358/openscience-3763), Shanmugam and associates described an interesting phenotype of Drosophila mutants missing either 7SK or the Larp7 protein, which is another component of the 7SK snRNP (Nguyen et al. 2012). In mutant larvae, neuromuscular junctions and animal locomotion are reported as defective. Unfortunately, it was not reported whether these earlier defects are associated with an adult wing posture defect similar to our *bin3* mutants. If the larval defects were shared with our *bin3* mutants, the specific function controlled by the Bin3-Box could still be related to 7SK function even though the level of 7SK remains normal in our Bin3-Box only mutants. Nevertheless, the possibility cannot be ruled out that Bin3’s role in maintaining normal wing posture is entirely independent of 7SK.

Our results showing the functional significance of the conserved Bin3-Box have important implications for the studies of MePCE. Prior approaches largely relied on RNAi knocking down of MePCE. As our results clearly show that such practice must have affected U6 steady state to a certain degree, which would have complicated the interpretation of results from any downstream assays. Our work thus offers a potential solution for this complication as we predict that disruption of the Bin3-Box alone in MePCE would generate separate-of-function mutations with little or no effect on U6 function. In addition, as neuromuscular development in flies and men share significant similarity, it might not be unexpected that other human patients suffering from neuromuscular dysfunctions similar to the reported case (Schneeberger et al. 2019) would harbor point mutations affecting other parts of the MePCE protein including its Bin3-Box.

## Materials and Methods

### Drosophila stocks and crosses

Flies were raised on standard corn meal food at 25°C. Unless otherwise noted, the *w^1118^* stock was used as the wildtype control. The chromosomal deficiency of the *amus* region *Df(3R)BSC47* was obtained from the Bloomington Stock Center (Stock # BL7443). The attP landing site at 75A10 (Stock # BL9725) was chosen for site specific integration of various *amus* transgenes. These insertions were introduced onto an *amus*-mutant chromosome by meiotic recombination. The actin5C Gal4 driver was obtained from Bloomington (Stock # BL4414).

For CRISPR/Cas9 mediated mutagenesis, an *U6*-driven sgRNA expression plasmid was injected into the embryos from a stock carrying the *vasa*-driven *Cas9* gene inserted on the *X* chromosome (Stock # BL51323). The survived G0 flies were mated to flies with balancers, and mutations were screened from F1 progenies with PCR and sequencing. The sgRNA targeted sequence for *amus* mutagenesis is 5’-GACATACGCCTGGACGTGCT**TGG** with the PAM sequence in bold. The sgRNA targeted sequences for *bin3* mutagenesis are 5’-GCAGCGAATTAAGGTTGAG**TGG** and 5’-GTTGCCGCCCAGCAGAAACT**TGG** with the PAM sequences in bold. All mutations were verified by PCR amplification followed by sequencing.

### Transgenic constructs

A 2.8kb genomic fragment (nt5732732-nt5735532 of chromosome *3R* in release dm6), containing about one kb upstream and one kb downstream of the *amus* coding region, was PCR amplified from wildtype genomic DNA and cloned into the vector pUAST-attB, and used as a rescuing transgene for *amus* mutations. Based on this transgene, a fragment encoding a GFP tag was inserted just upstream of the STOP codon of *amus* using bacterial recombineering (Zhang et al. 2014). Also based on the genomic transgene, the point mutations of Y83F, Y84A and K201A were individually introduced by site-directed mutagenesis. All DNA constructs were verified by sequencing.

An *amus* full length cDNA was reversed transcribed from wildtype mRNA and cloned into pUAST-attB. This *UAS*-driven transgene was combined with actin5C-Gal4 for Amus production. A cDNA clone encoding the first 76 amino acid residues of Amus and the last 275 residues of MePCE was synthesized using codons optimized for expression in Drosophila by Sangon Biotech in China. Point mutations disrupting critical residues for MePCE’s methylase activity were introduced into the above cDNA clone by site-directed mutagenesis. All of the above cDNAs were cloned into pUAST-attB for Gal4-driven production of the hybrid proteins. All DNA constructs were verified by sequencing.

### Cytology, Western blotting and FISH

Cellular localization of Amus-GFP was performed as followed: various tissues are dissected in PBS, fixed for 20 minutes in a fixative of 3.7% formaldehyde in PBS, washed three times for 15 minutes each with PBS plus 0.1% Triton T-100. After standard DAPI staining, tissues were washed then mounted in VECTASHIELD, and observed with a Zeiss LSM880 confocal microscope.

FISH analyses of U6 on testes and ovaries were performed with a protocol by Nizami et al. (2015). Cy3-labelled oligos were synthesized by Sangon Biotech in China, and used as probes, with their sequences listed in Table S2.

Amus full length protein tagged with 6XHis at its N-terminus was expressed as inclusion bodies and purified from bacteria using a standard IPTG-based induction and Nickel resin-based purification method. This full-length protein was used as antigen for animal immunization. Sera were used in Western blot at a dilution of 1:1000 for the detection of overexpressed Amus and Amus-MePCE proteins in extracts from adult flies or transfected S2 cells. An anti-MePCE antibody was purchased from Bethyl Laboratories Inc., and used at a dilution of 1:1000 on Western blots.

### Northern Blot and RNA-IP

Total RNA was extracted with a standard TRIzol reagent-based method. For Northern Blot analyses, 1μg of total RNA was denatured at 80°C for 3 min and separated on 5-8% 8M Urea-polyacrylamide gels in TBE buffer. RNA was transferred to a nylon membrane in TBE buffer at 20V for 30 min, dried and UV cross-linked. The membrane was pre-hybridized in hybridization buffer (25% Formamide, 4XSSC, 50mM NaH2PO4/Na2HPO4 Buffer pH 7.0, 1mM EDTA, 5% dextran sulfate, 0.5% SDS, 5XDenhard’s solution) at 42°C for 2h and hybridized with 0.1μg /ml of probes in hybridization buffer at 42°C for 16h. The membrane was washed sequentially with 2XSSC (with 0.1% SDS) for 5 min at room temperature, 0.2XSSC (with 0.1% SDS) for 5 min at room temperature, 0.1XSSC (with 0.1% SDS) for 15 min at 60°C, twice for each wash. The probes were oligonucleotides labeled with biotin, synthesized by Sangon Biotech in China. The signals were generated with streptavidin conjugated HRP, and quantified by Image lab software 5.2 (Biorad). Primer sequences are listed in Table S2.

RNA-IP was performed according to a published protocol (Gagliardi and Matarazzo 2016). Briefly, embryos collected overnight were bleach dechorionated, washed twice with cold PBS supplemented with Roche Protease Inhibitors, and pelleted by centrifugation. The pellet was suspended in lysis buffer (100 mM KCl, 5 mM MgCl_2_, 10 mM HEPES-NaOH pH 7, 0.5% NP-40, 1 mM DTT), supplemented with Protease Inhibitor cocktails and 400 U/ml of RNaseOUT from Invitrogen. The suspension was sonicated in a cooled sonicator (Sonics) followed by centrifugation for 15 min at 13,000 rpm. The supernatant was incubated overnight at 4°C with an anti-GFP nanobody coupled on agarose beads. The materials precipitated with the beads were washed twice with cold PBS and divided into two parts. One part was used for protein extraction with an elution buffer (50 mM Tris-HCl, pH 7.0, 5 mM EDTA, 10 mM DTT, 2% SDS) followed by Western blot detection of Amus. The other part was treated with Proteinase K at 55°C. RNA was extracted using TRIzol reagent followed by Northern blot detection of RNAs.

### PCR related experiments

For RT-PCR, total RNA was extract from 10 to 15 adult flies using TRIzol reagent. RT was carried out with the PrimeScript Kit from TaKaRa. Quantitative RT-PCR reactions were carried out on a Real-time PCR machine (QuantStudio 5 by Applied Biosystems), according to a protocol from the Bimake SYBY Green Master Mix qPCR Kit (B21402). Three biological replicates were performed for each sample. Primers are listed in Table S2.

qPCR for analyzing the expression from telomeric transposons was performed as described by Cui et al. 2021.

### Cell culture experiments

siRNA against *mepce* and control siRNAs were synthesized by GenePharma in China, with sequences listed in Table S2. siRNA reagents, used at a final concentration of 100 nM, were transfected using Lipofectamine RNAiMAX from Invitrogen as followed: a starting culture of HEK293 or HeLa cells at about 30% confluency was cultured for 24hr, followed by a first siRNA transfection. After an additional 24hr of culturing, a second siRNA treatment was administered. After the second transfection, cells were cultured for an additional 24 hr when they were divided into two parts. One part was used for protein extraction, followed by Western blot detection of MePCE with an antibody from Bethyl Laboratories Inc, (A304-185A). The other part was used for RNA extraction followed by Northern blot detection of RNAs. Probes for Northern blot are listed in Table S2.

Drosophila S2 cells were transfected using the FuGENE HD Transfection Reagent with two plasmids: one carrying an actin5C-driven Gal4 gene and the other carrying a UAS-driven *amus-mepce* expression construct. A mixture of 100 ng of actin5C driver plasmid and 400 ng of UAS expression plasmid was transfected. Transfected cells were cultured for 36 hours, harvested and divided into two parts. One part was used for protein detection by Western blotting, and the other used for RNA detection by Northern blotting.

### Protein sequence analyses

Homologs of Bin3/MePCE/Amus were retrieved from databases via Blast searches, and sequences are listed in Table S1. Phylogenetic trees in Marz et al. (2009) were used as a guide in our analyses. Sequence alignments were performed by Clustal Omega, the degree of homology was calculated by “Ident and Sim”. Motif analyses of the Bin3-Box was performed by the “GGSEQLOGO” R package. Protein phosphorylation predictions were made with “NetPhos3.1” and “PhosphoSVM”.

## Acknowledgements

We thank Dr. Steven Hanes of the SUNY-Upstate Medical University for sharing the *bin3^2-7^* and *bin3^4-7^* mutant flies. We thank Mr. Xionghong Tan for assistance in Drosophila cell culture experiments.

## Data availability statement

Drosophila stocks and plasmids are available upon request to YSR. The authors affirm that all data necessary for confirming the conclusions of the article are present within the article, figures, and tables.

## Conflicts of interest

None declared.

**Note**: While we prepare our manuscript, Palumbo and Hanes reported results from characterizing Drosophila *bin3* mutants (https://doi.org/10.1101/2023.06.01.543302) in which they observed a similar wing posture phenotype and identified a MSM motif similar to our Bin3-Box.

## Supplemental Materials

**Figure S1.**
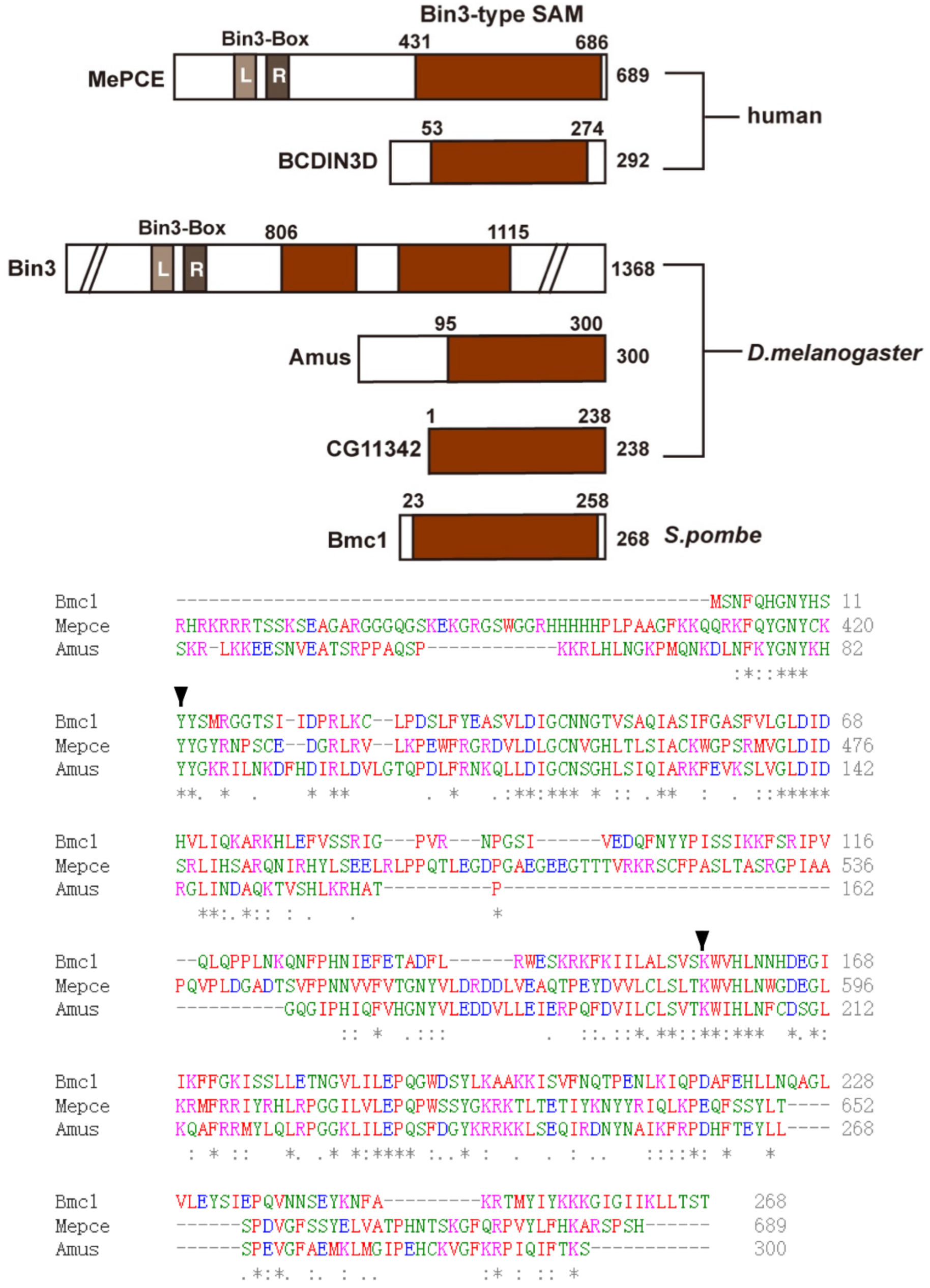
The Amus protein and its relationship with other Bin3-like methyltransferases. At the top is a schematic representation of domain structures of Bin3-related methyltransferases. The methyltransferase domain is highlighted in red and the range in amino acid is indicated on top of the diagram. The approximate position of the Bin3-Box is indicated for MePCE and Bin3. At the bottom are the alignments of the methylase domains from human MePCE, Drosophila Amus and S. Pombe Bmc1. Amino acid residues critical for the methylase activity (Y83/Y421 and K201/K485) are indicated by arrowheads, which have been individually mutated in this study.

**Figure S2.**
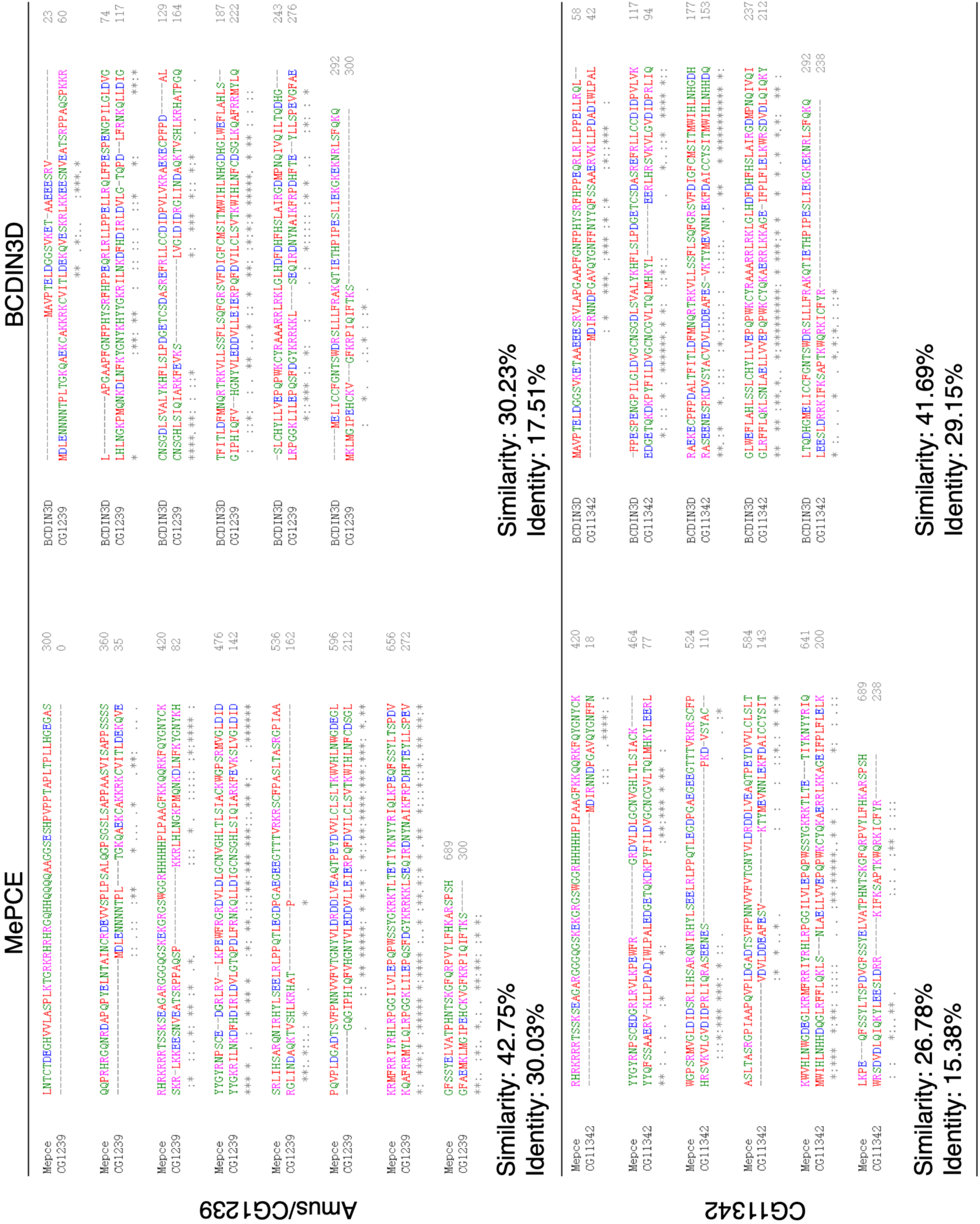
Amus is more related to MePCE than BCDIN3D. Pairwise comparisons between mammalian and Drosophila Bin3-related proteins (MePCE, BCDIN3D, Amus, CG11342). The percentages of protein similarity and identity are shown for each pairwise comparison.

**Figure S3.**
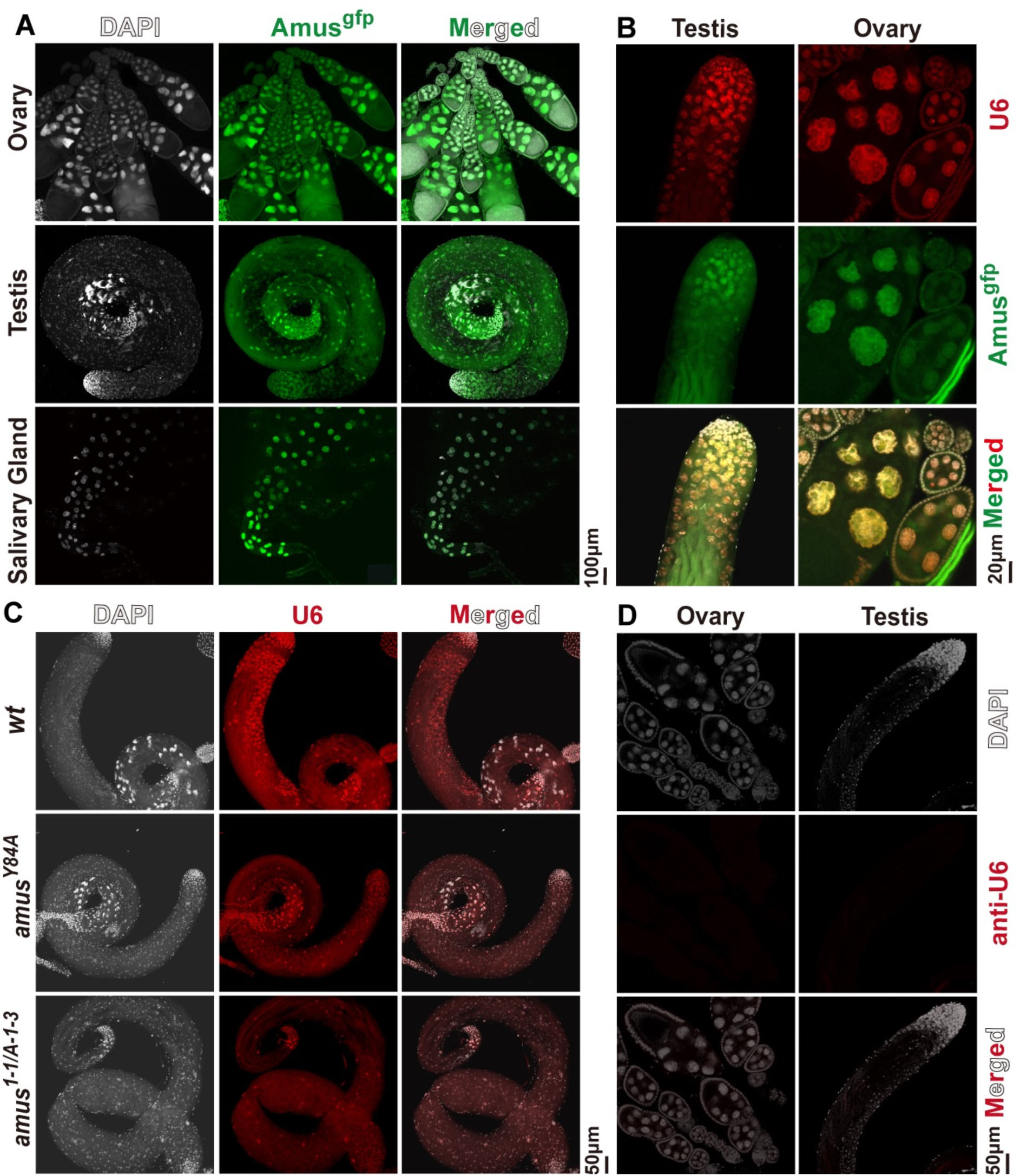
Nuclear localization of Amus and U6 snRNA. **A.** Images of lower magnification showing Amus^GFP^ localization in various tissues. Adult ovaries, testes and larval salivary glands were dissected, fixed and stained for DNA. DAPI signals are in white and GFP fluorescence signals in green. **B**. Amus^GFP^ co-localization with U6. Amus^GFP^ fluorescence signals are in green and U6 RNA-FISH signals in red. **C, D.** Images of lower magnification showing U6 localization in FISH experiments. A U6 sense probe (anti-U6) was used to confirm specificity of the signals from the anti-sense probe (U6).

**Figure S4.**
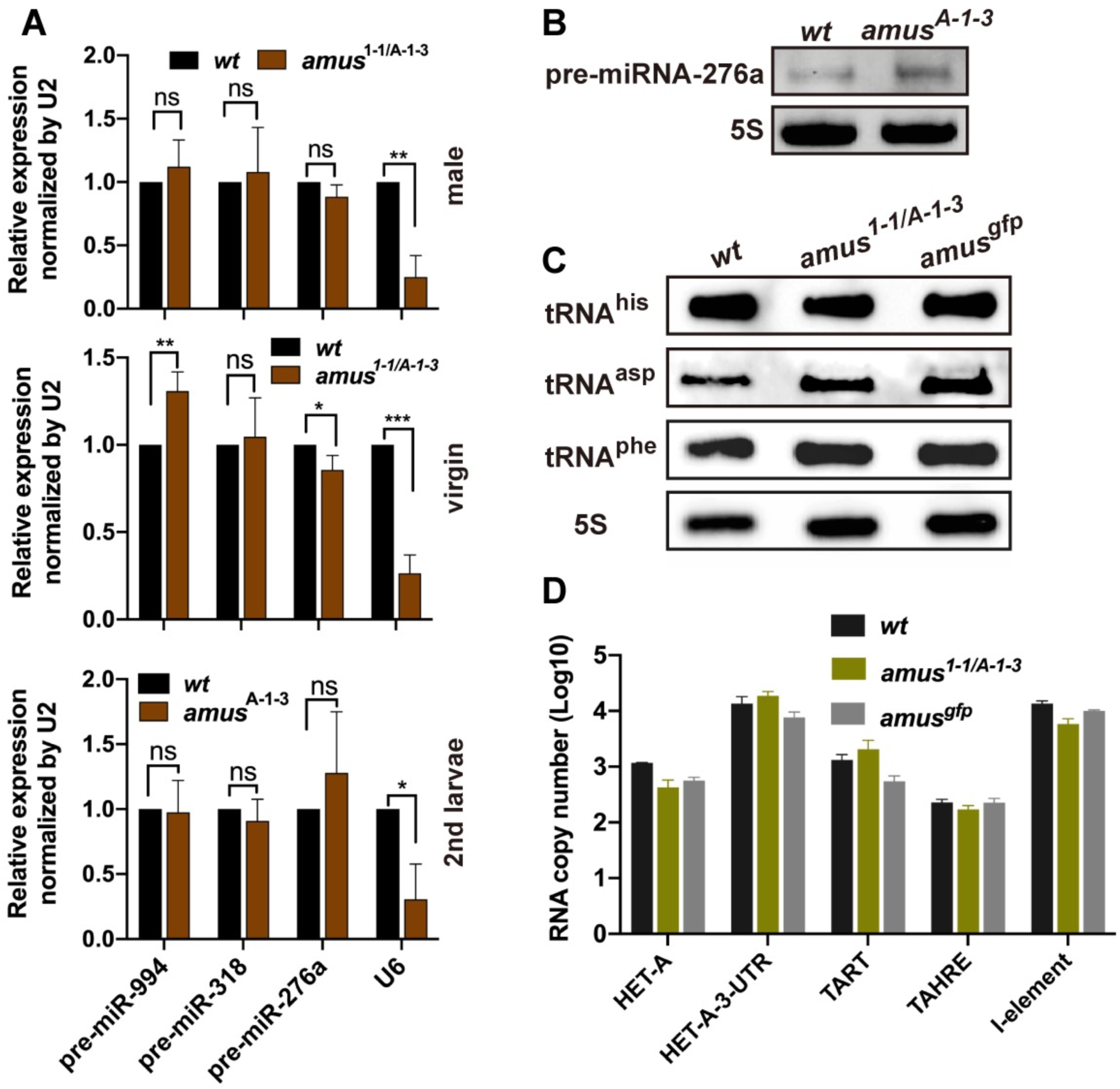
The effect of Amus loss on the levels of small and telomeric RNAs. **A**. Expression levels of representative pre-miRNAs measured by qPCR. Total RNAs were purified from adults, males or virgin females, and second instar larvae (mixed sex) of various genotypes. qPCR was performed in three replicates and normalized over the level of U2 snRNA. U6 snRNA was used as a positive control for the *amus* mutants. ***: *p*<0.001, **: *p*<0.01, *: *p*<0.05. **B**. Northern blot detection of pre-miRNA. RNA was purified from second instar larvae and probed with a probe against pre-miRNA-276a. The 5S rRNA was used as a loading control. **C**. Expression levels of representative tRNAs measured by Northern blotting. The names of the tRNAs are listed to the left and genotypes at the top of the images. The 5S rRNA was used as a loading control. **D**. Expression levels of telomeric transposons measured by qPCR. An absolute quantification method described in Cui et al. (2021) was used to measure *HeT-A*, *TART*, and *TAHRE* telomeric elements in adults of various genotypes. The non-telomeric *I* element was used as a control.

**Figure S5.**
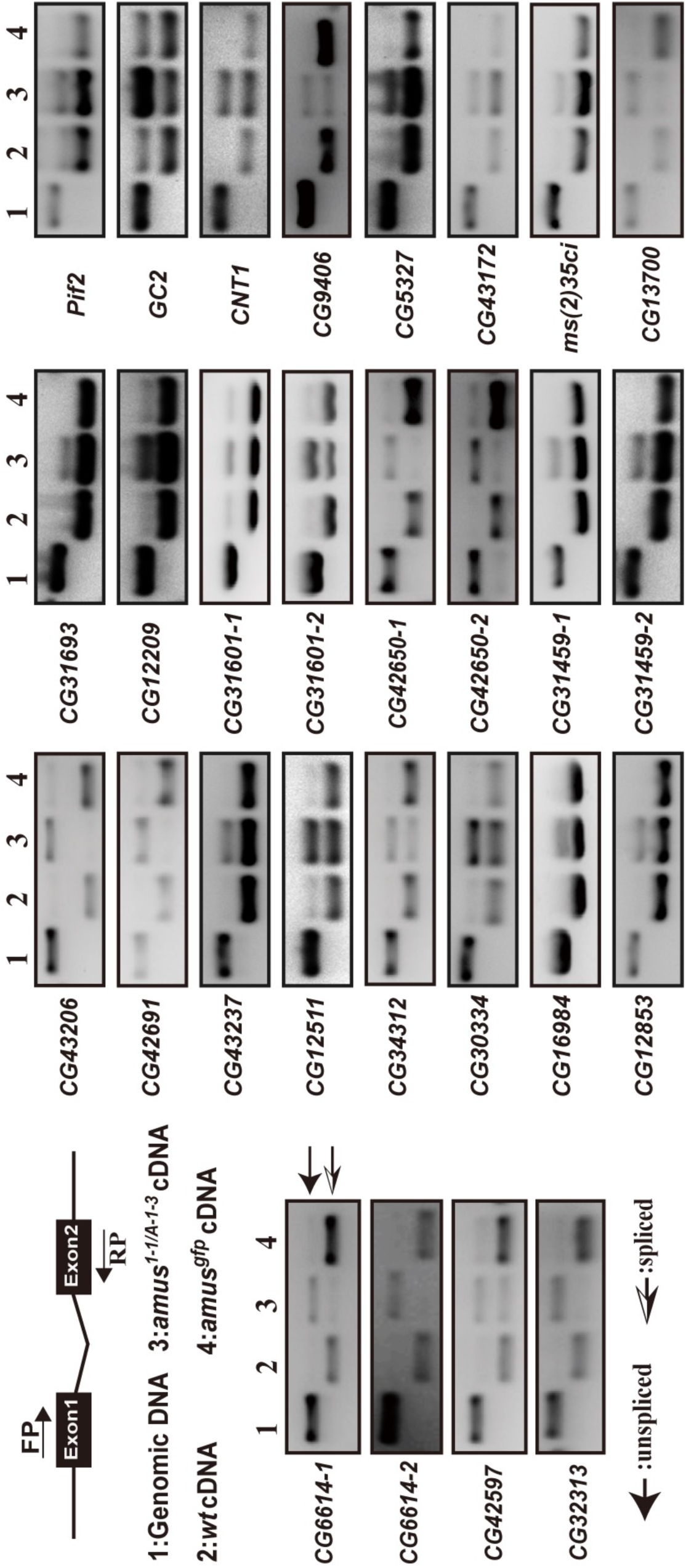
Intron retention events in *amus*-mutant testis. RT-PCR validation of intron retention events. The diagram at the top left depicts the approach showing a pair of PCR primers (FP and RP) spanning the intron of interest. Below the diagram, the four templates in the RT-PCR assay are listed. Gene names are listed to the left of the actual gel pictures from the RT-PCR assay with templates (1-4) at the top. The bands corresponding to products amplified from spliced and unspliced templates are indicated to the right. For some genes, retention events were detected for two introns.

**Figure S6.**
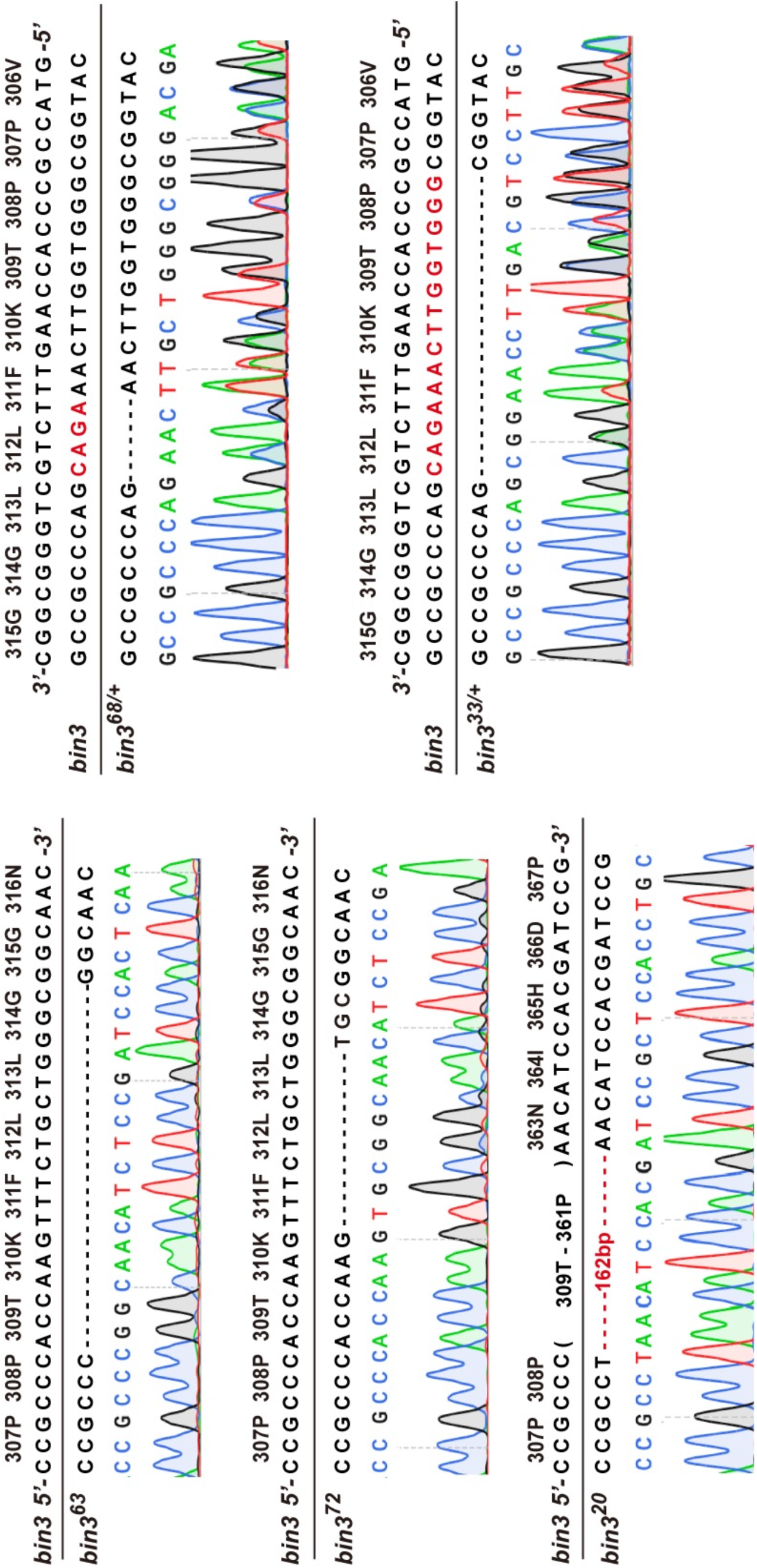
Mutations of *bin3*. Sequencing verifications of the newly generated *bin3* mutations. The original sequencing peaks were shown underneath the nucleotide sequences and amino acid sequences of the region. Deleted nucleotides were represented with “-”. For *bin3^68^* and *bin3^33^*, DNA from heterozygous adults were used in PCR amplification resulting in double peaks.

**Figure S7.**
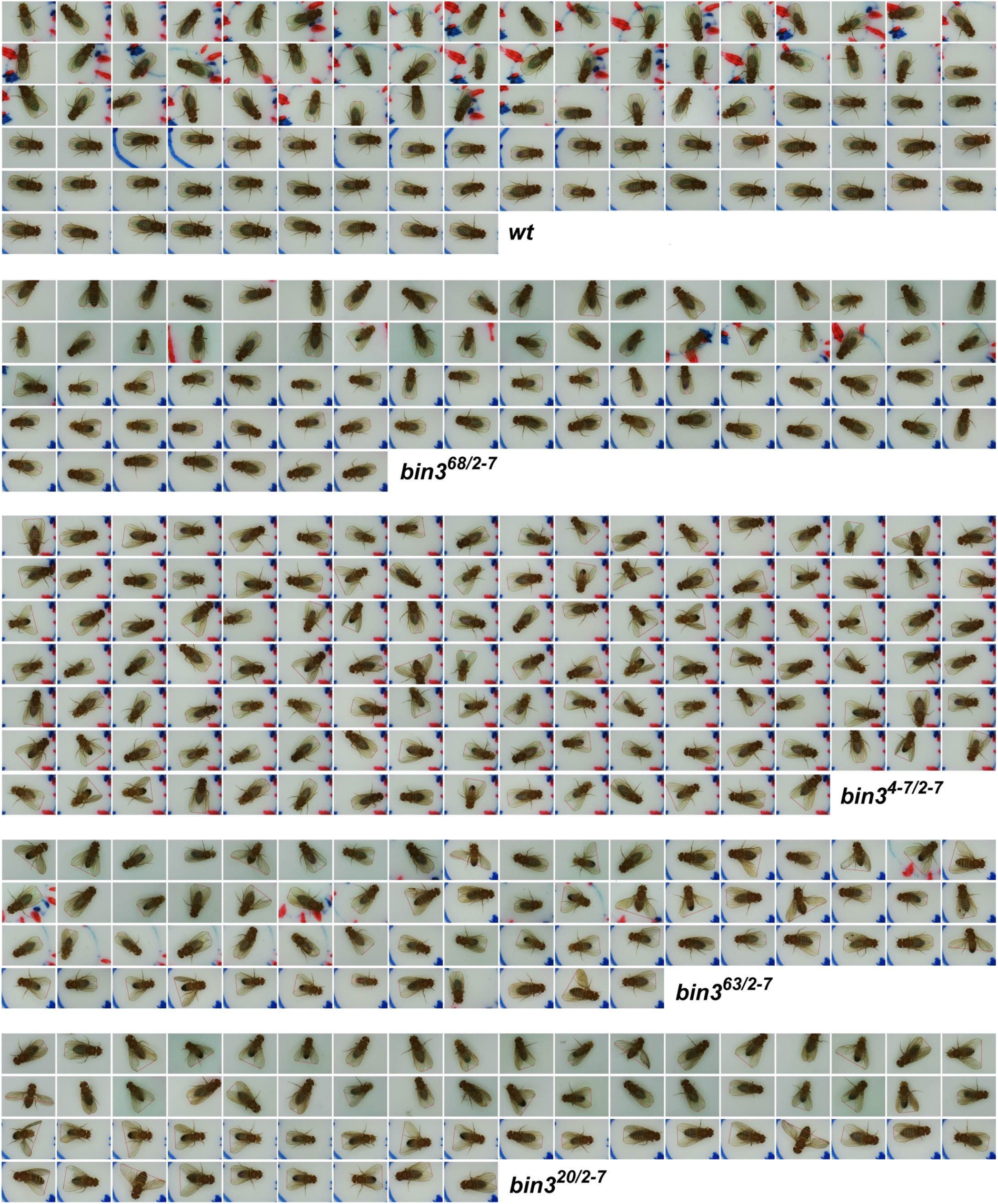
The wing posture defects in *bin3* mutants. Original pictures of the adults used for quantification of the wing posture defects.

**Table S1. Protein ID and sequences**

**Table S2. Sequences of oligos**

