## Supplementary material for "Drosophila Amus and Bin3 methylases functionally replace mammalian MePCE for capping and the stabilization of U6 and 7SK snRNAs": table s1

Table S1. Bin3/MePCE/Amus proteins

In organisms with only one MePCE-like protein, the name of MePCE will be used. In organisms with only one Amus-like protein, the name of Amus will be used, except for S. pombe. In organisms with two MePCE-like proteins, the name of Bin3 will be used for the one with the Bin3-Box and Amus used for the other.

>AAF51965.2 Amus/Dmel_CG1239 [Drosophila melanogaster]

MDLENNNNTPLTGKQAEKCAKKRKCVITLDEKQVESKRLKKEESNVEATSRPPAQSPKKRLHLNGKPMQNKDLNFKYGNYKHYYGKRILNKDFHDIRLDVLGTQPDLFRNKQLLDIGCNSGHLSIQIARKFEVKSLVGLDIDRGLINDAQKTVSHLKRHATPGQGIPHIQFVHGNYVLEDDVLLEIERPQFDVILCLSVTKWIHLNFCDSGLKQAFRRMYLQLRPGGKLILEPQSFDGYKRRKKLSEQIRDNYNAIKFRPDHFTEYLLSPEVGFAEMKLMGIPEHCKVGFKRPIQIFTKS

>AAF63187.1 Bin3 [Drosophila melanogaster]

MEKRLSDSPGDCRVTRSTMTPTLRLDQTSRQEPLPQQPDNGPAAAPGKSKSPTPLPGKSQAAQHHQFRAPQQQQGPKNRNKAWSKRQHKAGGKHNPSACVGASSGQAQSTGSSVIAATLLPTAASAHKADLENIQNIHNKNLTAGGGVNHHGNAGTAHHGGGGGAGAHHAAAGGHHHHHNTRLAQNAAAGGASGGGTIQMHKKMLRGHHHHVLCAGNNANHTCCLVTGCNGSSIGGVGVAGSGGATASAGGGGASCKEAQSCKDTSSLSGNSSIAGSAGAGNAVHYCCGRSKFFLPEKRLRKEVIVPPTKFLLGGNISDPLNLNSLQNENTSNASSTNNTPATTPRQSPITTPPKVEVIIPPNIHDPLHLLDPVDSMEYEKQLTSPMKRAGPGGGMLHHRQHHYRTRKNRKRRRFDSNNTSHAGDEGGVGSELTDEPPLPAATSSLAASPVAAPLNVGGSLLLSESAAPAPGETAEMGQQQEQAHVHSPQSASTTTTAAEMPTPTPTSAAAATATAEHKEQSAPAPTATSSPQRQQQHVAAAAEELPTPETSAAAETPAEEMLLSCSATSASLAVASTLAERRASRDLRLDLSSTCYGVGGTGLSFGGSISSSVGSSFGGGGRKRKISESSTSQKSKKFHRHDAMDKIVSPVVPQPGAWKRPPRILQPSGARKPSTRRSTSVSESELLSPVEEQPPKQLPLIGVEIPRDDTPDLPDHGLGSPLSTTSGATSHTAGEQDSLAGVDISMGDTLGSGVVGKAPLTSSLMLEPAKIPPIKMLPKFRADGLKYRYGNFDRYVDFRQMNEFRDVRLQVFQRHVELFENKDILDIGCNVGHMTITVARHLAPKTIVGIDIDRELVARARRNLSIFVRIPKEEKLLEVKAEPTVDAKANIAVKDETSGAAHKKTRRGKRRRKVHQGIHHHHHHHHDLEQLQQQQKLNSLLVKPHEFFPISFPLTYGRIPRILSSSKSPNMLGNKNQFPANVFFRHTNYVLKDESLMASDTQQYDLILCLSVTKWIHLNFGDNGLKMAFKRMFNQLRPGGKLILEAQNWASYKKKKNLTPEIYNNYKQIEFFPNKFHEYLLSSEVGFSHSYTLGVPRHMNKGFCRPIQLYAKGDYTPNHVRWSDAYYPQTPYEAYRGIYATLPVHRMGGGGSSAGGSNSGHAQMLHLSSSSRSQNYDTPHYAGSASGSASCRQTPMYQPTYNPLETDSYQPSYDMEYLNHMYVFASPLYQTVWSPPASLRKSSSHTPVFGSVRDAELDGDGSGGGGSGGGSYHRHVYPPNDDTCSPNANACNAFNSIRDADTDDSNQLPGGSRRHVYATNCGESSSSPQVNHHDAVGEFVDGLMDDEQKSSTGGGTGGAAYCDLSDA

>AAF47892.1 uncharacterized protein Dmel_CG11342 [Drosophila melanogaster]

MDIRNNDPGAVQYGNFFNYYQFSSAAERVKLLPDADIWLPALEDGETQKDKPYFILDVGCNCGVLTQLMHKYLEERLHRSVKVLGVDIDPRLIQRASEENESPKDVSYACVDVLDDEAFESVKTYMEVNNLEKFDAICCYSITMWIHLNHHDQGLRFFLQKLSNLAELLVVEPQPWKCYQKAERRLKKAGEIFPLFLELKWRSDVDLQIQKYLEESLDRRKIFKSAPTKWQRKICFYR

>XP_002408372.2 MePCE [Ixodes scapularis]

METAVEVPSLHVDVSSKRAGVFQRSTCRSASTDSQARPKTLKPANNSSRASPLVRKPPPREETASNQNRPPIGGKFVHGGGGNKRRQSFSVAADYKQPFNKRRRRGDLFTPPTKFLLGGNINDPLNLASFADDEVNKKANEVTPKSSPIPTPKHRTQVEVLIPPNINDPLNLNSGEDIEFNLISPRSRKRNRRRSKRKSRSEDDGPEQADASAPAAEVSTDADASARVASSSSTTASTTTTSRKASSVSAGAQPSFSKEGADKIVSPVVPQGSPTKFNRRKSSAAGGGASRDVRRSQHQGGRRAKQVYFREKDAHFQYGNYNRYYGYRNNGGQEDPRLKVMQASWLDGKDVLDIGCNVGHFTLSLARGFSPRKVIGLDIDGRLIKAARRNVRHYLTAALAGEMRFPVSMAICHGPIAAAALPGRESEPVTFPNNVFFVQGNYILESDVLLEAQREEFDTILCLSLTKWVHLNWGDEGVRRLFKRMFRQLRPGGRMLLEAQPFHSYVKKKKLTETTYKHFHDIRLRPEQFNEYLLSPEVGFASCELIDTPSHASKGFQRPIYLFTKAESAEEQCDAAQSECGSPTKAGCEKEGERDSGSEKPEREPPTEKRK

>XP_005178003.1 Bin3 [Musca domestica]

MDTVSNFDNPGDCQSKNLSPRVGNENNSPARHVANLVDHKRVEGATTAAIVDMEGKPTSASMPLATNKLKSQQQQQQHHQFRAPQQFVKHKKASSSTSTPSATTTATVTATANASSLNNNKKNKCWNKRQNRTAKHNTGSSNSSSSGGGGGSCGPQTAATTNNPSSPPNAQSLAEEKPGLTKADVENIQNLKNQGHNYYHNYPHPHYSSNHHGPGGGGGGGGIGNMPPNDNRLGLSLSSADKLGYGHGHLKTQQQPLHKKPLPSHRGGGGGGGGGGGGAAGGGGGGHNNHNHHHHHILCAGSAHNTMGSGVGNNAGSGTGVNANNLSCCHINGCGSHMGKDNQALNNNSNPNNSSASIQFCCIRSKFFLPDKRPRKGNFIPPTKFLLGGNISDPLNLSSLQNETGDTSIKITPATTPRQSPITTPPKVEVIIPPNIHDPLHLLDPVDSTEYEKQLTSPMKRAVVGGVGGGILGLPSAGVGGAIILGLRGAQNSKAHKHRHRKNRKPKRKRYDSCNSTTSTLGDALDDITPSALINASSSTEKGLLSVSLEAENKSQDQNNEGGVSSSGGDDDDNDRPERNLCSPQEQLSLSHNTADTKVANIDVDTTSQQTKSPTPTPVTQLAQEPMDSS

GAGMVGSVSTTTTAISFGEFRERACRDLRLDLISTTGSNSCGNSVVSSTGCGGGGGSRKRKISESNNSQKSKKYFRNDSMDKIVSPVVPQPGAWKRPPRTTASGARKSSNRRSRSISESNTADLSPVEENAKALVEEIPRDDTPDLMKVDSTSEFSALLGSPLSTASGGTSINAGEGDNMADLLNETGDASNALKLAENVKKEMDSMTVSTIKSIKPKFLPDGSKYRYGNYDRYAGLRHLNEFMDIRLQVFMQYPELFKNKDILDIGCNVGHMTISVARKLAPKSILGIDIDKELIARARRNLAMYVRIPVETWKKENVEVPEADVKNKDKYVKLKAGRRRRKKKKNRTDISQQRNNMNWGHFQHHHLMHYPHHHHHHHHHHHHHHHHHPHGAATAEQVGSKLDATEFFPISFPLCYGGITQSSAKMASPQNLESQSPASTTTSVSTTTKSTPGHPASGGGVGSGAKQQNSQNSSTNVRTQMSNLKNEFPQNVFFRQTNYILKDENLLANDTQQYDLILCLSVTKWIHLNFGDAGLKMAFKRMFNQLRPGGKLILEAQNWASYKKKKNLTEEIYNNYNNIEFFPNKFHEYLLSSEVGFSHSYTLGIPKHLSKGFSRPIQLYAKGDFTPSHIRWSDAYYPQTPFEAYRGIYATMPPRPTGMQSSYPLLSSTRSQNYDTPQYTGGGSGPYSCRQTPMHQPSTSQYYNPLESDSYQPSYDMEYLNHMYVFASPLYQTVWSPPTSMRKSSSHTPVFGSVRDAELEGDGNAMPGGSYHRHVYPPNDEISSPNANSGNAFNSIRDPDTDDSNQQRPHVYATNCGESSSSPHPHGGMNPSDTEAVDDSLILDEESRTNCCDLSEA

>XP_001656911.2 Bin3 [Aedes aegypti]

MEGQTLVLPERGGGGRDEGGGVGTEELRAFASVAEQGETAAEKTQLCPPRTRSQGGSNKKQQNKHFSRQQCKSKKWKLNKFRRNQKDGIGGGKKKDQAKGLDCGLTGGKGDEKQNPLGVNTECKQQQGKSKHSNKNNNSLMQARTKRLQSVSKFFLPDKRPRKDCIIPPTKFLLGGNISDPLNLNSLQNESENAVTPKSSPIPTPPHYKNKIEVIIPPNINDPLHLLDPVDSVEYEMQLCSPMKKKQRARNKRKRKSKSEKNNQSLDSSAGIAAGGQLSTSISEAEATTAEEAQMEAPGSTGKNVEHSLEGGNTGDRDKATRCLRLDLEVCRKRRISESTCSSKNKVRRMDSMDKIVSPVIPQPGAWKRGHWPAMPVGGRNRTRTQSCTSNSEEPPGKQP

VLEGEVNTETPQTTEPDIKVNNVEQPKPEPSESETQAKPKPHEDSKYHYGNYDRYYGYRNLNEFIDVRLKVFLRNPYLFRDKDVLDIGCNVGLMTIAVAKMLHTKSITGIDIDEKLIAKARRNLTTHVRIQPGHAKHGGETKLSGEKRKRRISRKLTTGEEKSLVKEEEAPHEEKPAEVPDTAESMPKPETSSAKKGQHPHRRKFPGRHSHGKHHMHNHHHNHQRKDPRKGNSEFYPISFPLSLGNFGSKACLLDMGAKDHKFPNNVIFKTMNYVLKDESLINYDTQQYDLILCLSVTKWIHLNFGDNGLKMAFKRMFNHLRPGGKLILEAQNWASYKKKKKLTDTTCENYKSIEFFPNKFHDYLLSPEIGFSHSYPLGIPRHLSKGFRRPIQLYVKGDFTPSQAQWSDTYHPQTPYVNHHNIYTDITTPTQNPCIWGSTTPYQPSYSTRCTPSHSFMATSPYYNPRQTDSYQPSYDAGNRRGSYCFASPLYSTTWSPPSDLRHRRSASNTPNSASIRSAENDEQFAQQQHRPHVYPPMEVFPLESTAVRSVAPGSASVRQTDSDSEPSSNSASQTPTRSQYDVDQSSNSPSTTHSQPDK

>XP_004930468.Bin3 [Bombyx mori]

MGSPEEKITACEAVNENEARDTIDKSELNTSDITESAANESNEQILNDANASKDSDTPGAGISKKAKKRSRHGKKGHTPSKDNVDRNVVRRVHYTGTKFKQGRKRAQSFPSISKFFLPHKRPRKETFIPPTKFLLGGNISDPLNLNSLQDEDVNRAMNAVTPESSPLPTPPRHKAKIDVIIPPNIRDPLNLMEPLDNEEYEKRLEMQQRKPKKKTKVRRRTTDNSSPTKEVSEEEVIESIKVPQTPEASMSAQNLRKRSCSDSENSSQKGKDKKLRRMDSLDKIVSPVVPQPGAWMRARPQARPAPPERPVSPHRTKLKSNSEGEQQLPTFHPKNSRYQYGNYDRYYGYRNLNAMMDIRLQVFEMHRHLFQNKDVLDIGCNVGHISIAVARTFGARSVVGIDIDSVLIGRARKNLQSFVAVPPPEPPKTDCDKKTDVEDKKIDCNKCHQEKCDDFCEQKEKADDPKKTIPGRKLKKPRKNRKSFYKHEQGQSFFPMSMSMLYGPIRDLLPELLTFPHNVTFKHGNYVPREDLAVGSGESPQFDLILCLSTTKWMHLNWGDAGLKRAFRRMFADLRPGGKLVLEAQNWASYKKKKRLTPTIYENFNSIELFPIKFRDYLLSPEVGFSKCCILGVPQHASKGFRRPIQLYIKGDFTPSKARWSDAYVPSSHHRPEAEQPRRTVYAPVEPAYRPSDDAPSCGAATPYSPNLDCCTDDSPFYNPRSDSYAPSYQPPRPTVYATLPSPLGSASPAQSPAASPAPHASPAPDPLTARGFPEPRPHDD

>XP_002129700.1 MePCE [Ciona intestinalis]

MATAVTMSPSNCDAMKQQKPDIVSSVQVPSGNTGQSSSGNTGKLEIKCKPIMSAFPVRGKRFHRKRRHSTSEEFPGSINAAKSQNAKHRRQGASHHKKEIAGDIILPTNFLLGGNIRDPLNLNSMLDEKVNRALNAVTPSSSPLPPRNSTISIVIPNDMTDPLGLNAAPTEEVAENSSLNQPSVQKKERERSNSKSRNRKRRKKSLVERTLPDQEDTLQDEGKNEPDSITGVLKLDNKCMKVKKNKPSFLVLDGNCDQNSLSVHVPQTSAVTDPIVSPVIPQKTNRNVVFSPKSRKRKASASIHAKNIAQLNMDVVPGTDLLVVASETKETLEKKGAKKSLYSHENESTSEKTCVQYSAVCSPSQNSQKTEKKGTNVKSAKPSFKKNNARFQYGNYSQYYGYRNPKKFQDCRMTCLKREWFKEKDVLDVGCNVGHLTLLIARDYSPRKIVGIDIDGSLINIARTNIRHYATASVRPPSQLASSFPDSMPKMYGPLVAPPVETGKDDSAEFPNNVLFFEGNYVPEVDEILRFQEPEYDVIMCMSVTKWMHMNWGDEGLKKTFERFFKQLRAGGKLIIEPQEWKSYGKKKKLTERIYKNYNSIELKPAMFPDYLIKVVGFESVEYLTLTTQNRSEGFKRPIQLYRKPE

>XP_031755492.1 MePCE [Xenopus tropicalis]

MQVLLEMAATTENLQLPGAAGSTEGEATTFLAPPPAPPPPKAELQPKATKEGAEQQAAPRGPRNGFQGKRRNSFNVGFKHPVPFKRRRRVNSDCDPVLPSNFLLGGNIFDPLNLNSLLDEDVNRTLNAETPKSSPLPSRNRDPVEILIPKDITDPLCLNASSGELLLSPLKSRKRHRHRHHPPTDAPSKAPGGPAQEAEPAPEEPQPYELNTAINCRDEVVSLPALESDPAAPSAGASAISASRHRKRRRTSSKSEGKHGSPERSKSLLAEGAALRPHCPKSRAKGKTRFQYGNYCRYYGYRNPSQSEDPRLRALKPEWFRGKAVLDIGCNVGHVTLCVAKNMGPSRVVGLDIDRSLITAARQNIRHYLSGETKRDGGFPAGLVATRGPIAAPSLPPTGGEGNGFPHNVVFVQGNYVVEREELVEAQQSEYDVILCLSVTKWVHLNWGDEGLKRMFRRMFRHLNPGGTLILEPQPWASYGKRKKLSEAIYKNYSKLAFRPEQFTQYLVSPEVGFSSYELIGTPPTHSKGFQRPLYAFHKSAPCAGTEKLPDPASPGAAPACGEGI

>NP_001122001.1 MePCE [Danio rerio]

MIEMSFDKETVLSCEGQTDPTSQEFNINDTSALHPPGNTTENNLSKDAGSEASLEKSATTDNNASSESAVLGAESSKQSHNGTKPQQQKVNKRRNTMSAGFKHPGYGKRRRRANSESESVLPTNFLLGGNIFDPLNLNSLLDEEVNKALNAETPKSSPLPAKNRDPVEILIPRDITDPLNLNCLSTGDSLLVSPLKSGGGRRRHRNKHHGANTGRVSTEGIGLTESNGCAAVKDDGTSLVSMTNVHLESTIADQTLLDSTLESSNKSSLTSNDNEVKEKTVDCSTAHLTANPTNPPSSRQRKRRRAGSRTENSSSSSEVKRSGRHFQHNESQGGNSRSGVSDRQQPYPNQWRPQYSNNNQNPQQPKKKFQYGNYNKYYGYRNPGMSEDPRIRVMNPDWFRGKDVLDLGCNTGHLTLFIAKNWRPASIVGLDIDGSLIHAARQNIRHYLSEVQVQHSRRSGENTKADRGEVSGEEKDKDKTSLVMDEKKAKHVDEVNKCMDEGMEVNQEEKREADRGEIGDGASVDLPDGKHSFPVSLRISRGPIAGPPLPETNTHSLPPGDFPANVTFIKGNYVLESDVLLQTQREEYDVILCLSVTKWVHLNWGDAGLKRFFHRVYKHLRPGGLFILEPQPWSSYNKRKKLTEAICKNYHSIRLKPDQFSSFLTTEVGFSSYELIGTSQNYSKGFQRPISLYHKRPSSLK

>NP_062552.2 MePCE [Homo sapiens]

MIEMAAEKEPFLVPAPPPPLKDESGGGGGPTVPPHQEAASGELRGGTERGPGRCAPSAGSPAAAVGRESPGAAATSSSGPQAQQHRGGGPQAQSHGEARLSDPPGRAAPPDVGEERRGGGGTELGPPAPPRPRNGYQPHRPPGGGGGKRRNSCNVGGGGGGFKHPAFKRRRRVNSDCDSVLPSNFLLGGNIFDPLNLNSLLDEEVSRTLNAETPKSSPLPAKGRDPVEILIPKDITDPLSLNTCTDEGHVVLASPLKTGRKRHRHRGQHHQQQQAAGGSESHPVPPTAPLTPLLHGEGASQQPRHRGQNRDAPQPYELNTAINCRDEVVSPLPSALQGPSGSLSAPPAASVISAPPSSSSRHRKRRRTSSKSEAGARGGGQGSKEKGRGSWGGRHHHHHPLPAAGFKKQQRKFQYGNYCKYYGYRNPSCEDGRLRVLKPEWFRGRDVLDLGCNVGHLTLSIACKWGPSRMVGLDIDSRLIHSARQNIRHYLSEELRLPPQTLEGDPGAEGEEGTTTVRKRSCFPASLTASRGPIAAPQVPLDGADTSVFPNNVVFVTGNYVLDRDDLVEAQTPEYDVVLCLSLTKWVHLNWGDEGLKRMFRRIYRHLRPGGILVLEPQPWSSYGKRKTLTETIYKNYYRIQLKPEQFSSYLTSPDVGFSSYELVATPHNTSKGFQRPVYLFHKARSPSH

>AAH26876.1 MePCE [Mus musculus]

MIEMAAEKEPFLVPAPPPPLKDESGGGGGPEVQSHQEAASGELRDGTEHGPGPRAHSAGAAASGGGGPQAQAHGEPHGRAAAPADVGEERRGGGGTDLGPPAPPRPRNGYQPHRPPGGGGGKRRNSCNVGGGSGGSFKHPAFKRRRRVNSDCDSVLPSNFLLGGNIFDPLNLNSLLDEEVSRALNAETPKSSPLPAKGRDPVEILIPKDITDPLSLNTCTDEAHVVLASPLKIGRKRHRHRGPHHQQQQASGGNDSNAAVLPTDPLTPSLHGEGATQQQQNRGQNRDAPQPYELNTAINCRDEVVSPLPSALQGSSGSLSAPPAASVTSAPSTSSSSRHRKRRRTSSKSEAGARGGSQGSKEKGRGSGGGRHHHHPLPATGFKKQQLKFQYGNYCKYYGYRNPSCEDVRLRVLKPEWFQGRDVLDLGCNVGHLTLSIACKWGPARMVGLDIDPRLIHSARQNIRHYLSEELRLQAQTSEGDPGTEGEEGTITVRKRSCFPASLTASRGPIAAPQVPLDGADTSVFPNNVVFVTGNYVLDRDELVDAQRPEYDVVLCFSLTKWVHLNWGDEGLKRMFRRIYRHLRPGGILVLEPQPWSSYCKRKSLTETIYKNYFRIQLKPEQFSSYLTSPEVGFSSYELVATPNNTSRGFQRPVYLFHKARSPSH

>NP_496573.1 MePCE [Caenorhabditis elegans]

MSSHRRGSFRGRKRFYRDTFAPGGSKTDPLNIEIELTENPEEEKKRLGLLDPTTERSKKRKVEKDEKSEKPAENSPFKKNQYNSPRKDSRRPPQLSKEEKSAAENRKQKTEYFNKKYRYGNFDRYYGIRLNPGESDKRLSVFQKDWFEHKQALDIGCNAGFLTLSIAKDFSPRRIIGIDIDEHLIGVARKNIRHYCDHETEVSGKFPASFGVQFGTVSQRNEAPRSFSTKFPDNIWFKKENYVLESDEMLDMIQPEFDVILALSITKWIHLNWGDDGMRRFFRRAYAQLHPGGRLIIEPQAFDSYKKRAKMSEELKANYSKIEFKPEDFEMWLIETVGFESVEKLGVVGAKSKGFERPIDVYLKPLHPKTDAIPLGYI

>XP_008121627.2 MePCE [Anolis carolinensis]

MIEMATDKEAFRVPAPPPPLPKPPPKGAAACGAGDLPGKAAGKGGGGAGKGQGARGGGDGGLSAPASASADPQRGARPLGRRATAWPSSAGLAMASRPHSRLHRHSPQRPNKRRNSCNVGGGGGFKHSAFKRRRRVNSECDPVAPSEFLLGGNIFDSLNLNSSLGEEVNQGSQRRDLQVVLAAGQRARSRVGAAPRRRTKGPAPPPPPASACQKPCNKPCNKPCNGATPQPYELNTIINCRDEVVSPLLPAGHGEPPPGHQQVSGGCVTPSCASVASSASASSRHHRKRRRTCSKSEGAATRHASTEQPPRSSPERAPRSAPRSRHPPPASARRQPRHKFQYGNYCKYYGYRNPDCEDVRLRVMKPEWFQGKEVLDVGCNVGHFTLSVAKKWGPARMVGLDIDGHLIHSARQNIRHYLSEELHQQKQQQSDEGPSPGARKTFPASLMASRGPIAAPRVPQEGTDASVFPNNVIFIKVRANYVLERDELLEAQRPEYDVILCLSLTKWVHLNWGDEGLKRLFKRIYRHLRPGGILVLEPQAWSSYKKRKNLTETISRHFNRIKLKPDQFPSYLTSSEVGFCSYELVAMPRNTPKGFQRPIYLFHKTRPGGH

>XP_002166936.3 Bin3 [Hydra vulgaris]

MEVTQKPKSPIPSPNNIRNLSSKIISPKLKFDFNYRLNTGVRRKCSEKKYHPLKMIFVHDPLNLDSLTSKSNGIESPNMRSPVPDRSCEFQPRLLQPVDLRDPLSLNCLDDTAQILTPLRKRRKRKRRFSAGDPYDTLSFSMTEISPKKSSFKFDTQKNNVCDDLLIRSDPSCVECKKEKTSFVKKDIYRSLTKPNLLCEKKKKKEKTFTYGNYIKSTKDDERLSCFSKQWFFNKNCLDIGCNNGKVTLEILKRFDPSFITGVDIDDRLIKLARSYARNDSYMPSRQRFPVSIKQTYGPMLNTRILQHGLSGNICFITQNYVPCELDDLSKIKEEYDTITCLSVTMWVQLNFGDSGLKRMFKKIFMQLKPGGKLLLEPQLLKSYKKKKNLSITILHHFKSIKLFPNDFVRYLLSKEIGFATSECLIYTHHKKTGGRPLFLLTKSSGSNIV

>XP_047122849.1 Amus [Hydra vulgaris]

MKRVHVEKDCKFKYGNYNRYYGYRNENTDVDKRISLLQKDWFEGKSCLDIGCNVGHITLYIAKFFEPKQIKGVDIDFNLIKAARANILHYIDNKNSESRKVKKSNEPNESNQKNSKINQIENESLIINGEQLNCCCFEKKDEQLLQLHEDVTKCKHDKNSSSNCIHNGNGNVQAEHLKREKKFPYNVTFITENFVPSSENLLKYTKEEYDIIICLSVTKWVQLNSGDEGLKLMFHKIFKLLNPGGKLLLEPQPYKSYKRRKNLSTEIRNNYDNIKLKPEDYINFLMSEVGFTSFTQLDTLKHDKQGFERPVYLLTKASTVL

>RDD42018.1 MePCE [Trichoplax sp. H2]

MEVNMSGATTWRQNFISETSPSHQERGKKANLSYKKTKKAKVRRSLSSEKSLLPSKPQDPTDPLNLRSLLDPEINNRLKADTPHVSPSHRRLANVKNVTNEDQILVPPQTSDPLMLSREDREYDKRVRSLKQHKKKRKLSKDNDDKNDLQTLKKIRQDKVELVETTLKEKNSKQKLFIYGNYNRYYGYRNQSTHDGRLDSWNHQWFQGKDCLDIGCNDGHLTIEIAKKFNPNTIVGIDIDASLITRARSNITRINSTKNHQFPISFGICHGPIVDTCNDDSTLYPRNILFKQENYIKESMESINEEKPLYDVILCLSLTKWIHLNWGDEGVKKLFKRIFMNLRHDGILILEPQSIESYKKKKRLSATFRKNFDEIKLMPDQFNDYLLSEEIGFSSCQTLPIPNDAVKVYDAIRDISNHPMMIIYLFVIVFWTHNVSIRKKLMLSTSRWIPS

>NP_568752.1 Amus [Arabidopsis thaliana]

MGRDNDQKKNKKKRNRSNENEKSVEKVVANEEKVPTQQKQKQQQGQQGNCNQSKKKKNQEVYPFGNYRNYYGYRISNDTDEDPRLKVLKKEWFEGKDCLDIGCNSGIMTIHIAKKFGCRSILGVDIDTSRIEDAHWHLRKFVRMQNSTKPSEKKSSSEGADGVHGSKEPSVSLSNGEAKADSAETKDLSQIVSFQKENFVQTRNLDDNRYDTILCLSVTKWVHLNWGDDGLITLFSKIWRLLQPGGIFVMEPQPWKSYENNRRVSETTAMNYRTIVLRPDRFQEILLDKIGFRTVEDLTSSLSGASKGFDRQILAFQK

>CAD8097073.1 Amus [Paramecium primaurelia]

MNEQFRKFLLNQEKSDVQQKKSDSAPQIRLKLQSKRKTNVDLGLETFVQKKVKQDEKEDKKQPNLMEKLQMNPIDLVKSTKQYSYGNYKKYYHLRLQQKWEDPRLTILDSIYFENKSILDIGCNDGTLTLLIALKHYPKLIRGIDIDYTLINKAIEQMVHLDDQQKKIQKQEFKPIIEDLPVSFHKYMEQPMSKAIEQQYIHQTIEDMNKQNEHNNNSILFNQIAKNNIFPHNVYFRVQNIIGNKKYDEKYDTILCLSITKWIHLNFGDIGIKRLFKTISNFLNVGGHLILEPQEWKSYKKKKYYSSEFKQNYKEIQLKPQDFSKVLEKEYNFKLIQQINPDDESAIKKSKSTFRRPILIFEKQNEI

>NP_596220.1 Bmc1 [Schizosaccharomyces pombe]

MSNFQHGNYHSYYSMRGGTSIIDPRLKCLPDSLFYEASVLDIGCNNGTVSAQIASIFGASFVLGLDIDHVLIQKARKHLEFVSSRIGPVRNPGSIVEDQFNYYPISSIKKFSRIPVQLQPPLNKQNFPHNIEFETADFLRWESKRKFKIILALSVSKWVHLNNHDEGIIKFFGKISSLLETNGVLILEPQGWDSYLKAAKKISVFNQTPENLKIQPDAFEHLLNQAGLVLEYSIEPQVNNSEYKNFAKRTMYIYKKKGIGIIKLLTST
