## Supplementary material for "Drosophila Amus and Bin3 methylases functionally replace mammalian MePCE for capping and the stabilization of U6 and 7SK snRNAs": table s2

**Table S2. List of oligos**

**1. List of probes used for Northern blot and FISH**

**Fly-Probe Name	Probe Sequence**

| U6 FISH (1-46) probe | 5'Cy3-CTCTGTATCGTTCCAATTTTAGTATATGTTCTGCCGAAGCAAGAAC |
| --- | --- |
| U6 FISH (62-103) probe | 5'Cy3- ATGTGGAACGCTTCACGATTTTGCGTGTCATCCTTGCGCAGG |
| U6 Biotin probe (3-49) | 5’ Biotin-CTTCTCTGTATCGTTCCAATTTTAGTATATGTTCTGCCGAAGCAAGA |
| U6 Biotin probe(68-107) | 5’ Biotin- AAAAATGTGGAACGCTTCACGATTTTGCGTGTCATCCTTG |
| 5S Biotin probe(1-40) | 5’ Biotin- GGACGAGAACCGATGTATTCAGCGTGGTATGGTCGTTGGC |
| 7SK Biotin probe(229-268) | 5’ Biotin- ATACGGCTTCCGGTCGAGACCAGAATCCGCGGGGTGGTGG |
| U2 Biotin probe(75-119) | 5’ Biotin- TCCGTCTGATTCCAAAAATCAGTTTAACATTTGTTGTCCTCCAAT |
| U1 Biotin probe (71-101) | 5’ Biotin- GAATAATCGCAGAGGTCAACTCAGCCGAGGT |
| tRNA^his^ Biotin probe(1-31) | 5’ Biotin- CGTGGGGTCCTAACCACTAGACGATCACGGC |
| tRNA^asp^ Biotin probe(1-36) | 5’ Biotin- GACAGGCGGGGATACTAACCACTATACTATCGAGGA |
| tRNA^phe^ Biotin probe (1-31) | 5’ Biotin- TCTAACGCTCTCCCAACTGAGCTATTTCGGC |
| pre-miR-276a Biotin probe(1-40) | 5’ Biotin- AACGTAGGAACTCTATACCTCGCTGATGGCAAAAACCAGG |

**Human-Probe and siRNA	Probe Sequence**

| 5S Biotin probe (71-110) | 5’ Biotin- CACCCGGTATTCCCAGGCGGTCTCCCATCCAAGTACTAA |
| --- | --- |
| 7SK Biotin probe (147-191) | 5’ Biotin- ACCCTTGACCGAAGACCGGTCCTCCTCTATCGGGGATGGTCGTCC |
| 7SK Biotin probe (12-52) | 5’ Biotin- CCCTGGCGATCAATGGGGTGACAGATGTCGCAGCCAGATCG |
| U1 Biotin probe (111-152) | 5’ Biotin- CGAACGCAGTCCCCCACTACCACAAATTATGCAGTCGAGTTT |
| U2 Biotin probe (116-155) | 5’ Biotin- GGTCGATGCGTGGAGTGGACGGAGCAAGCTCCTATTCCAT |
| U6 Biotin probe (60-99) | 5’ Biotin- TATGGAACGCTTCACGAATTTGCGTGTCATCCTTGCGC |
| U6 Biotin probe (1-49) | 5’ Biotin-CTTCTCTGTATCGTTCCAATTTTAGTATATGTGCTGCCGAAGCGAGCAC |
| si-mepce-1 | 5' GGGUAAUUAUGUGCUGGAUTT |
| si-mepce-2 | 5' GCAUGUUUCGCCGGAUCUATT |
| si-mepce-3 | 5' CCGCUCAGUCUCAAUACUUTT |
| si-mepce-4 | 5' GGCGGAGAAGGAGCCGUUUTT |

**2. List of primers for plasmid construction and** **verification**

**Primer Name	Primer Sequence**

| *amus*-gDNA-rescue-FP | tctgaatagggaattgggaattcgttaacagatctgcggccgccacctaggcaacttaatctaattat |
| --- | --- |
| *amus-*gDNA-rescue-RP | acagaagtaaggttccttcacaaagatcctctagaggtacctgaggaaaacaaccacgataa |
| *amus-*c-gfp-FP | AGCACTGCAAAGTAGGATTCAAGCGACCGATCCAAATTTTTACAAAGTCG |
|  | GGCGGAGGGCGCGCCCTGTGGAACACCTACAT |
| *amus-*c-gfp-RP | CCTTGAATTTTCAGATAATTTTCATTTCTTGATTTTCTTTTTATTTCCTA |
|  | CTTGTACAGCTCGTCCATGCCGAGA |
| *amus-gDNA-FP* | ATAAGAATGCGGCCGCCACCTAGGCAACTTAATCTAATTAT |
| *amus-gDNA-RP* | CGGGGTACCTGAGGAAAACAACCACGATAA |
| *amus^Y84A^*-gDNA-rescue-FP | TATAAGCACTACGCAGGCAAGCGCAT |
| *amus^Y84A^*-gDNA-rescue-RP | ATGCGCTTGCCTGCGTAGTGCTTATA |
| *amus^K201A^*-gDNA-rescue-FP | GCCTGTCTGTCACTGCATGGATCCACCTT |
| *amus^K201A^*-gDNA-rescue-RP | AAGGTGGATCCATGCAGTGACAGACAGGC |
| *amus^Y83F^*-gDNA-rescue-FP | TATAAGCACTTTTACGGCAAGCGCAT |
| *amus^Y83F^*-gDNA-rescue-RP | ATGCGCTTGCCGTAAAAGTGCTTATA |
| *amus*-cDNA-FP | ATAAGAATGCGGCCGCATGGATTTGGAGAACAACAACAATACGC |
| *amus*-cDNA-RP | CGGGGTACCCTACGACTTTGTAAAAATTTGGATCGGTCG |
| *amus-mepce*-cDNA-rescue-FP | agggaattgggaattcgttaacagatctgcggccgc |
| *amus-mepce*-cDNA-rescue-RP | aagtaaggttccttcacaaagatcctctagaggtacc |
| *amus-mepce^Y83F^*-cDNA-rescue-FP | GCAACTATTGCAAGTTTTACGGTTATCGAA |
| *amus-mepce^Y83F^*-cDNA-rescue-RP | TTCGATAACCGTAAAACTTGCAATAGTTGC |
| *amus-mepce^K585A^*-cDNA-rescue-FP | TGTCCCTCACAGCATGGGTCCATCTGA |
| *amus-mepce^K585A^*-cDNA-rescue-RP | TCAGATGGACCCATGCTGTGAGGGACA |
| *amus^A-1-3^* *wt*-FP | TCACGACATACGCCTGGA |
| *amus^A-1-3^ mut*-FP | TCACGACATACGCTTGGC |
| *bin3-*verification-FP | CAACGCAGTCCACTATTGCT |
| *bin3-*verification-RP | GGTGTTGTTGGAGTCAAAGC |

**3. List of primers used for qPCR and RT-PCR**

**Primer Name	Primer Sequence**

| *rp49*-qPCR-FP | ATGACCATCCGCCCAGCATAC |
| --- | --- |
| *rp49*-qPCR-RP | GCTTAGCATATCGATCCGACTGG |
| *rpl32*-qPCR-FP | GCCGCTTCAAGGGACAGTATCTG |
| *rpl32*-qPCR-RP | AAACGCGGTTCTGCATGAG |
| *pre-miR-318*-qPCR-FP | ATGGGATACACACAGTTCAGTTTTGT |
| *pre-miR-318*-qPCR-RP | CTCATGAGATAAACAAAGCCCAGTGA |
| *pre-miR-994*-qPCR-FP | CGAGTTATCTAAGGAAATAGTAGCCGTG |
| *pre-miR-994*-qPCR-RP | GAGCTATCTAAAAGAAACAGCAACTGTG |
| *pre-miR-276a*-qPCR-FP | GCCATCAGCGAGGTATAGAGTTC |
| *pre-miR-276a*-qPCR-RP | TTGGTCTTCCAAGAGCACGGTA |
| *fly-U6 snRNA*-qPCR-FP | GTTCTTGCTTCGGCAGAACATATAC |
| *fly-U6 snRNA*-qPCR-RP | TGTGGAACGCTTCACGATTTTGCG |
| *fly-5S rRNA*-qPCR-FP | GCCAACGACCATACCACGCTGAATA |
| *fly-5S rRNA*-qPCR-RP | AAGTTGTGGACGAGGCCAACAAC |
| *fly-U1 snRNA*-qPCR-FP | ATACTTACCTGGCGTAGAGGTTAACC |
| *fly-U1 snRNA*-qPCR-RP | CCCGGCTAACAAAAATTACACGC |
| *fly-U2 snRNA*-qPCR-FP | ATCGCTTCTCGGCCTTATGG |
| *fly-U2 snRNA*-qPCR-RP | CAACCCGTGACAGAGGTGGA |
| *CG42691*-FP14 | ATGTGCTGCAATGTGAGTCG |
| *CG42691*-RP378 | GCGCCTACAATCAGAACAAC |
| *CG43237*-FP25 | ATGAGCCCGGATATATAAT |
| *CG43237*-RP282 | CTACACCACATCATTTGACC |
| *CG12511*-FP118 | TGAGTGACTTACCAGTGGTG |
| *CG12511*-RP387 | ATGTCTTGAGCACATCCAGC |
| *CG34312*-FP1681 | CCATTAGGAAGTGCTGTTGC |
| *CG34312*-RP1951 | AACACTAGGTGGGCAAAGTG |
| *CG30334*-FP88 | TACCGAATAGAGGAGCTCAC |
| *CG30334*-RP337 | CACAGAAGGTGGTGCCAAAG |
| *CG12209*-FP548 | TTGGTATGTTCGCTCTGGTG |
| *CG12209*-RP815 | GAGTACGGCATATCGATCAG |
| *CG42650*-1-FP273 | GTTTGATGACACAGAGGACG |
| *CG42650*-1-RP541 | AAGCGGCTTCCAGAGCATTC |
| *CG42650*-2-FP481 | CAGCTGAAGAATGAAATCTC |
| *CG42650*-2-RP714 | AAGTCGTATCGGGCCTGTTC |
| *CG31459*-1-FP556 | TGATGTTGAATGCCTGGCAG |
| *CG31459*-1-RP798 | ACACATGAGCACCACGATTG |
| *CG31459*-2-FP721 | TCATCGGTCCTTTGCCATTC |
| *CG31459*-2-RP979 | AGTCGCGAATGTCAATCAGC |
| *GC2*-FP83 | ATGTTGGAACAAGTTGAGCA |
| *GC2*-RP328 | TGTACATGCGCTCTCCATTG |
| *CG9406*-FP95 | GTATATACCCGCGATGGAAG |
| *CG9406*-RP355 | GATAGTCGACGGAATCGGTG |
| *CG5327*-FP104 | GCGAAATGATCGAGGACATG |
| *CG5327*-RP354 | TATTCTGGAATTGGCCCAGC |
| *CG43127*-FP118 | TATATGATGCGACGTCGTCG |
| *CG43172*-RP378 | AGATGCCAGCTATCATGAGC |
| *ms(2)35ci*-ND-FP606 | CCAATCACATGGTGATTGCC |
| *ms(2)35ci*-ND-RP787 | TCCGATTGGTAGTAGCTCAG |
| *CG13700*-ND-FP311 | AAGGTTCACACGTCATCGTC |
| *CG13700*-ND-RP550 | TCGGTAGTCCATTGTCTGAC |
| CG6614-1-FP1307 | TGTTGCTGCAACTGTACCAG |
| CG6614-1-RP1597 | GTTCATTGCGCTCGCAAATG |
| CG6614-2-FP1806 | GCTGGGGAACATCTGTAATC |
| CG6614-2-RP2105 | TTGCTGACCAGTCGCATTTC |
| CG42597-FP64 | ATATTTACAAGACCGCGCAG |
| CG42597-RP314 | ATTACCTCTGTGTCGCACAG |
| CG12853-FP22 | CGAGCAACATCGATTTTGGCTG |
| CG12853-RP309 | TTTGCCTTTGCCCTTCTTGG |
| CG31639-FP51 | GCTAGTGAACGATCAGTTGG |
| CG31639-RP293 | ATAGGGACCGCATCCATAAC |
| Pif2-FP45 | ATAGTCACACGGACACACAC |
| Pif2-RP263 | ATTGCAGCACGGTGGACAAC |
| CNT1-FP1471 | TTATCGCAGCCAGCGTAATG |
| CNT1-RP1748 | GGAAGAACACAATGGCTAGG |
